# Characterization of DNA extrusion scenarios underlying aggregate Hi-C patterns by a new computational framework

**DOI:** 10.1101/2021.06.28.450073

**Authors:** Xizhe Zhang, Grigoriy Gogoshin, Sergio Branciamore

## Abstract

Aggregate Hi-C maps centered on several genetic elements show significant DNA extrusion features. However, these features are missing from the component Hi-C images. To resolve this discrepancy, we present a new computational framework that reconstructs DNA extrusion scenarios (DESs) and explains the enhancement of DNA extrusion features by aggregation. We showed that the reconstructed DES is essential to understand the site interactions of the relevant genetic elements. For example, CCCTC-binding factor (CTCF) binding elements (CBE) with a specific motif direction are only affected by DNA extrusion activities initiated from one side, manifested as a directional bias. About half of promoters search for enhancers in a specific direction, as indicated by the corresponding DES. Moreover, since DNA extrusion does not pause at promoter or enhancer elements, direct promoter-enhancer contacts are not long-lasting; rather, promoters and enhancers are brought in proximity by continuous, dynamic DNA extrusion.

## 2 Introduction

Comprehensive genome-wide measurement of chromatin interactions using Hi-C has identified several significant features of three-dimensional chromatin structure, such as compartments, topological associated domains (TADs), and loops [1][2][3]. Several hypotheses have been proposed to explain how the Hi-C features are formed (reviewed by[4]). One model postulated by Riggs [5] and independently by Nasmyth [6] suggests that these features are formed by a procedure called DNA extrusion, during which a loop extrusion factor (LEF), which includes cohesion, binds to DNA and, while staying in position, reels in DNA to form a gradually increasing loop.

DNA extrusion will fold the chromosome into a tandem array of loop structures. It also suggests a mechanism regarding the proximity of two distal elements to each other. In this study, anchor refers to the part of LEF contacting DNA. Each LEF has two anchors indicating that DNA could be reeled bi-directionally during extrusion[7]. The information coded in the DNA sequences is transferred to LEF through the anchor interface, causing pause, stop, or other states changes of DNA extrusion.

On the other hand, the polymer interaction model describes the chromatin fiber as a series of monomers and simulates the monomer interactions based on a set of constraints, such as the distance, the excluded volume, and the rigidity of the DNA fiber. More recent modeling studies have recapitulated some Hi-C features, like TAD and loops, by combining these two earlier proposals[8][9][7][10].

Several lines of evidence have supported the DNA extrusion model. First, DNA extrusion is consistent with the convergent CTCF rule, which indicates that the upstream anchors and TADs are preferentially associated with forward CCCTC-binding consensus sequences (CTCF motif, which is asymmetrical). In contrast, the downstream anchors are associated with the reverse orientation of CTCF motifs[3][9][11]. Second, loop extrusion factors (LEF), which are responsible for reeling DNA to form loops, have been identified; these include in eukaryotes condensin, cohesin, and in bacteria the bacterial structural maintenance of chromosomes (SMC) complex[12][13]. Third, the process of LEF binding to DNA has been further revealed. It was shown that the loading of LEF onto the genome is dependent on the protein Nipbl, and the releasing of LEF from DNA is regulated by the protein Wapl[14][15][16]. Fourth, the movement of DNA predicted by the DNA extrusion model has been confirmed by the in vitro system[17][18].

Computational biology studies have provided additional new insights that corroborate the DNA extrusion model[8][10]. For example, these studies successfully reproduced Hi-C features, such as the loop corners and domain flames (domain strips, see Fig 1), by modeling DNA extrusion[10][19][20]. They also demonstrated that the heterogeneity of DNA extrusion could be simulated by experimenting with a number of specific features such as adjusting the velocity of anchor movement, the permeability of barriers, and the loading and releasing of LEF from the DNA.

**Figure 1:**
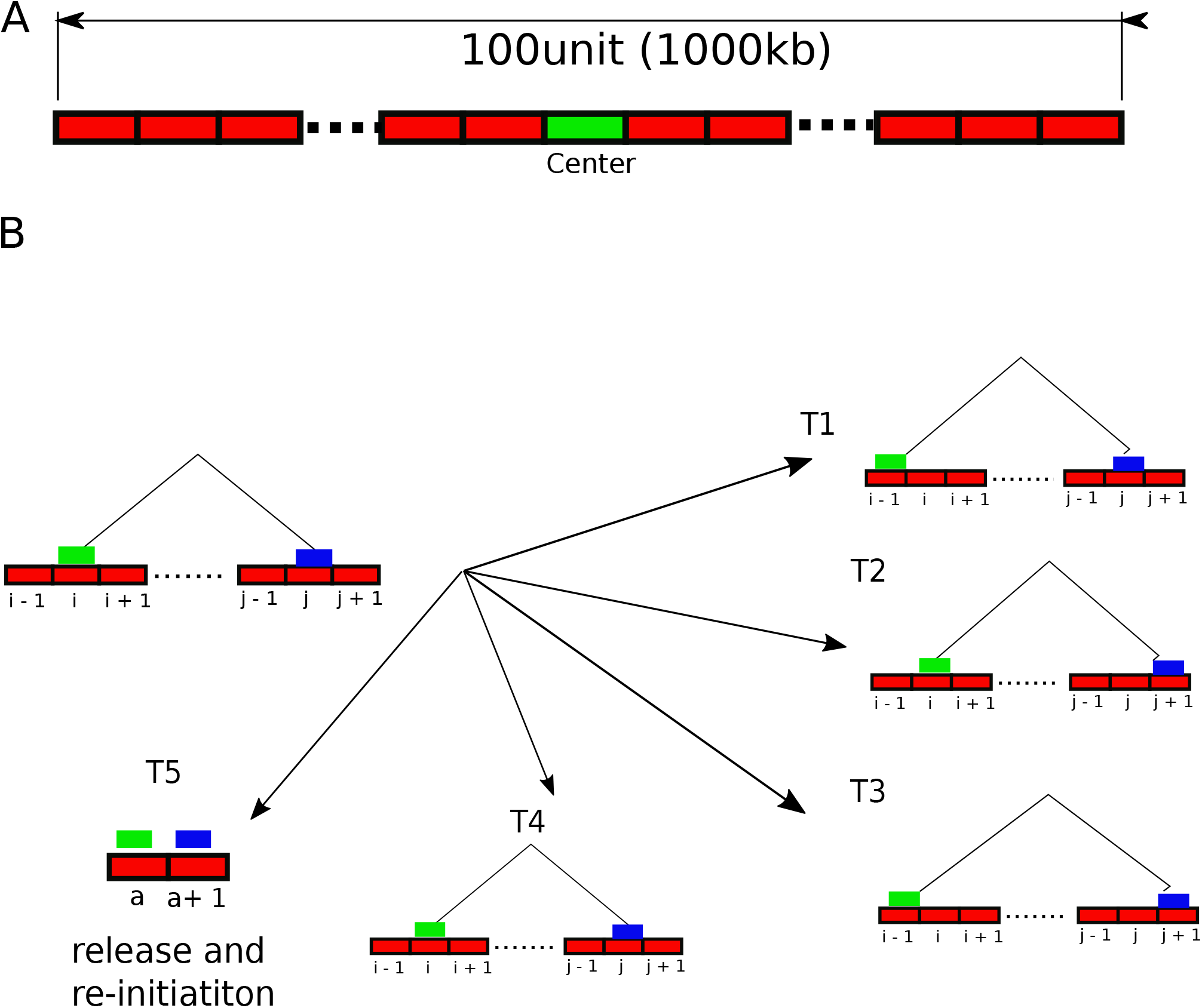
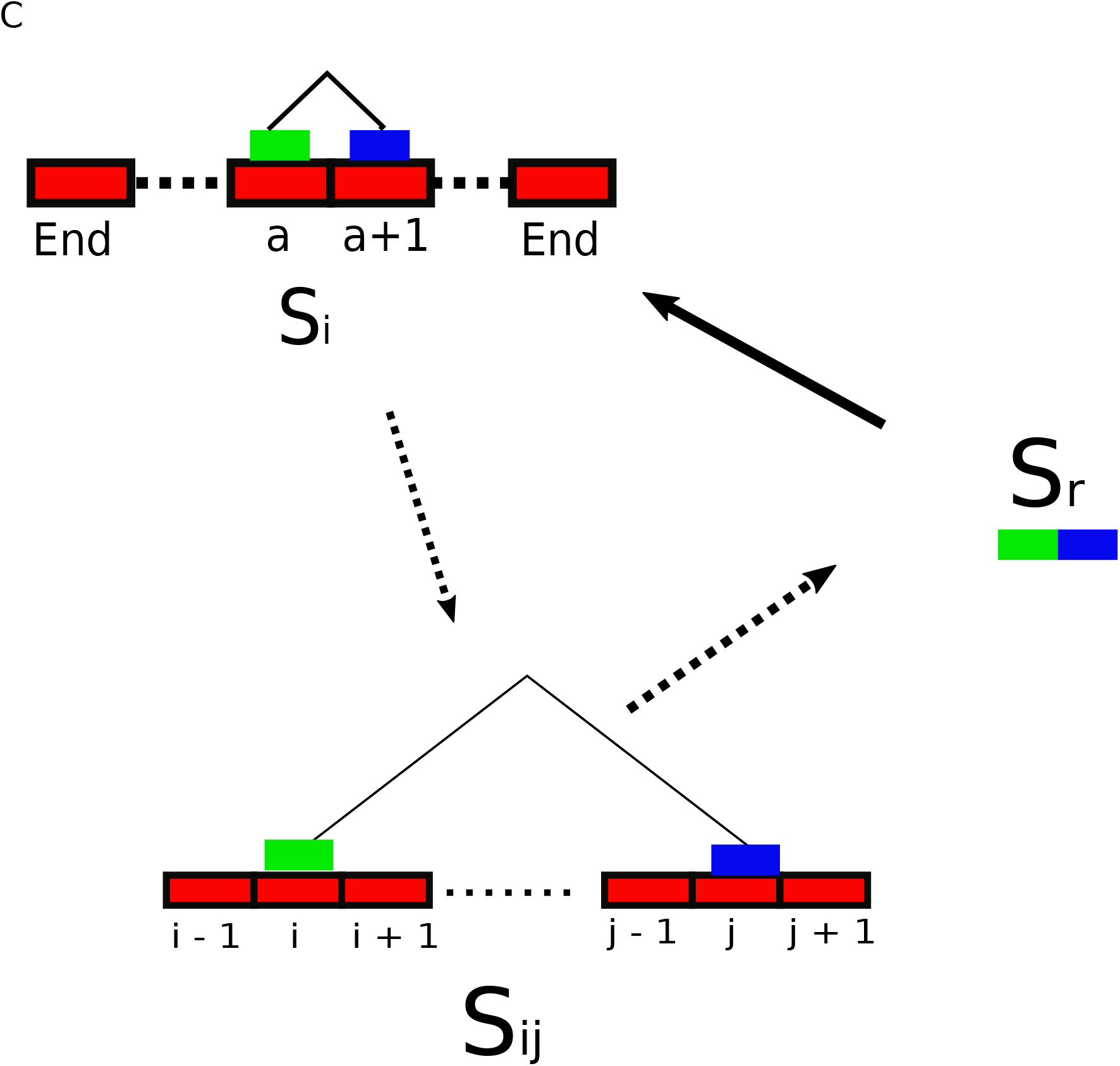
Modeling DNA extrusion (loop extrusion). (A) The DNA undergoing loop extrusion is defined as a one-dimensional lattice structure. Each unit of the lattice represents a 10kb genomic bin. (B) Shown the five transitions (T1-T5) from a loop configuration located at the *i*_*th*_ and *j*_*th*_ positions on the DNA lattice to the next configuration. (C) Shown are the loop configurations of DNA extrusion. The loop configuration *S*_*i*_ represents the initiation of DNA extrusion. *S*_*ij*_ represents a general loop configuration of which the anchors are located at the position i and j. *S*_*r*_ represents a special loop configuration in which LEF is released from DNA when one anchor moves out of the boundaries of the DNA lattice. The loop configuration after *S*_*r*_ is *S*_*i*_, which means a new round of DNA extrusion begins.

Due to the technical constraints, the captured ligation products reflecting the loop conformations formed by DNA extrusion are negligible compared to those formed by local polymer interactions. Similarly, compared with the polymer interaction model, the DNA extrusion model only explains a tiny fraction of the entire Hi-C data[10]. Therefore, a new method to identify and analyze DNA extrusion features is needed.

In this study, we found that aggregate Hi-C maps show significant DNA extrusion features, which could be modeled to reconstruct the DNA extrusion scenarios (DESs, See below). Our analysis of the DES associated with CTCF binding elements (CBEs) revealed the tendencies of these sites to interact with other regions with a strong directional bias[11][21]. We also demonstrated that tandem CBEs with the same motif orientation are associated with a dominant DNA extrusion event rather than multiple. In addition, we have two new discoveries pertaining to promoter enhancer interactions. First, about half of promoters search for enhancers in a specific orientation as a result of imbalanced directional DNA extrusion activities. Second, DNA extrusion does not stop or pause at promoters or enhancers, suggesting that these two genetic elements do not stably interact directly for a significant time but are brought into proximity by continuous, dyamic DNA extrusion.

## 3 Results

### 3.1 Modeling DNA extrusion by Markov Chain

We modeled the DNA extrusion with the following specifications. First, the DNA undergoing DNA extrusion was defined as a one-dimensional lattice structure composed of 100 units, and each indexed unit represents a 10kb genomic bin(Fig1a)[8]. Second, the DNA extrusion process was broken down into a series of loop configurations. The movement of loop anchors on the DNA lattice caused the transition from one configuration to another(Fig1b), if the DNA is considered as a fixed reference. A general loop configuration was represented by the positions of its left and right anchors on the DNA lattice. Third, there were two unique loop configurations. One was the loop configuration anchoring at two adjacent positions on the DNA lattice representing the loading of LEF to DNA(*S*_*i*_ in Fig1c). Another was the configuration representing the releasing of LEF from DNA when one anchor moves out of the boundary of the DNA lattice(*S*_*r*_ in Fig1c). Fourth, we assumed that the transitions depended entirely on the current loop configuration, such as interaction between loop anchor and a barrier element (BE). Finally, the loop size does not decrease. That is, the anchors will not change the movement direction during extrusion.

The above DNA extrusion model can be mathematically expressed as a Markov Chain (MC, see methods for details). Each event of the MC sequence represents a specific loop configuration during DNA extrusion. The whole MC sequence is considered to be a collection of DNA extrusion trajectories. Here, the DNA extrusion trajectory is defined as a set of sequential loop configurations from the binding of LEF to DNA (*S*_*i*_ in Fig1c) to the release of LEF from DNA (*S*_*r*_ in Fig1c). The state space of the MC corresponds to the entire loop configurations. Therefore, the simulated Hi-C image, corresponding to the equilibrium probability distribution of the entire states, is obtained by computing the eigenvector of the state-transition matrix.

Of note, we defined transition probabilities associated with a state (loop configuration) to be random, except for the states related to a BE or DNA extrusion starting point (DSP). Here, a DSP is defined as the DNA region where DNA extrusion is initiated. We calculated various DNA extrusion scenarios (DESs) using the MC model with defined BE, DSP, and the transition probability matrix. The defined conditions of BE and DSP are listed in table 2. Finally, our modeling did not consider the local DNA polymer interactions, but we show that this model is consistent with the aggregation process and sufficient to reproduce the significant features of the aggregate patterns.

### 3.2 Modeling CTCF aggregation patterns reveals that aggregation enhances DNA extrusion features

Aggregate Hi-C mapping or pile-up analysis has been widely used to enhance the Hi-C features associated with important regulatory elements such as enhancers, promoters, and transcription factor binding sites[22][13][23]. We generated an aggregate Hi-C map centered on the genome-wide CBEs in GM12878 cells (total 25894, see methods for details) and named the recognized pattern as the CTCF aggregate pattern (CAP, Fig2a). For this study, we divided CAP (and other aggregate patterns in this study) into three areas (Fig2b): two smaller triangle areas symmetrical to the center (named sub-domain A, SDA and sub-domain B, SDB) and the middle square area (called the insulated domain, ISD). The CAP shows the following features. First, the signals within SDA and SDB are higher than the signals within ISD, which identify clear boundaries between ISD and SDA or SDB (two red line segments, Fig2b). Second, there are no apparent sub-patterns found within SDA and SDB. Third, the signals are enriched within the boundaries, forming a structure similar to the domain strips[19]. Finally, the whole CAP pattern is nearly symmetrical to the center.

**Figure 2:**
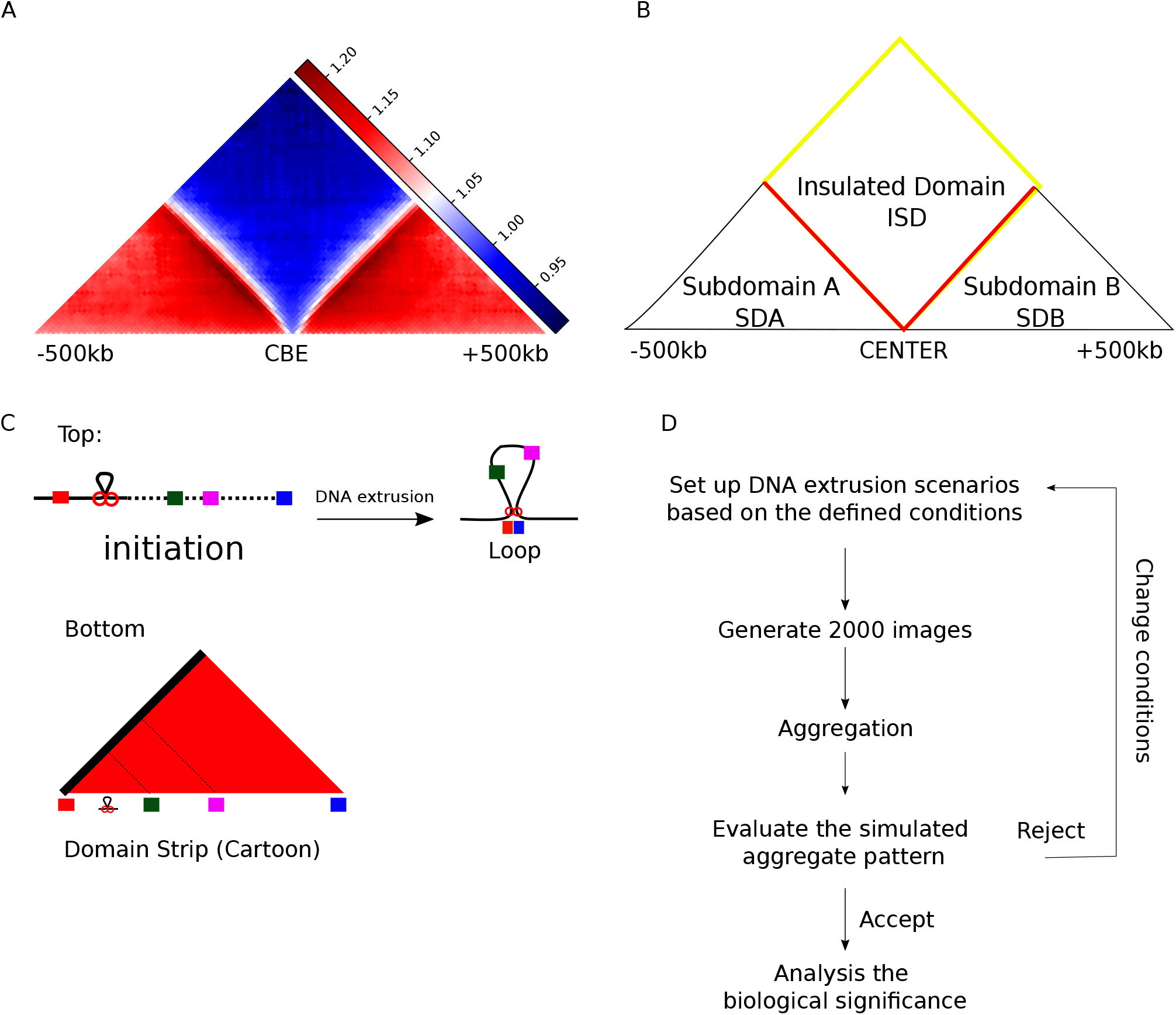
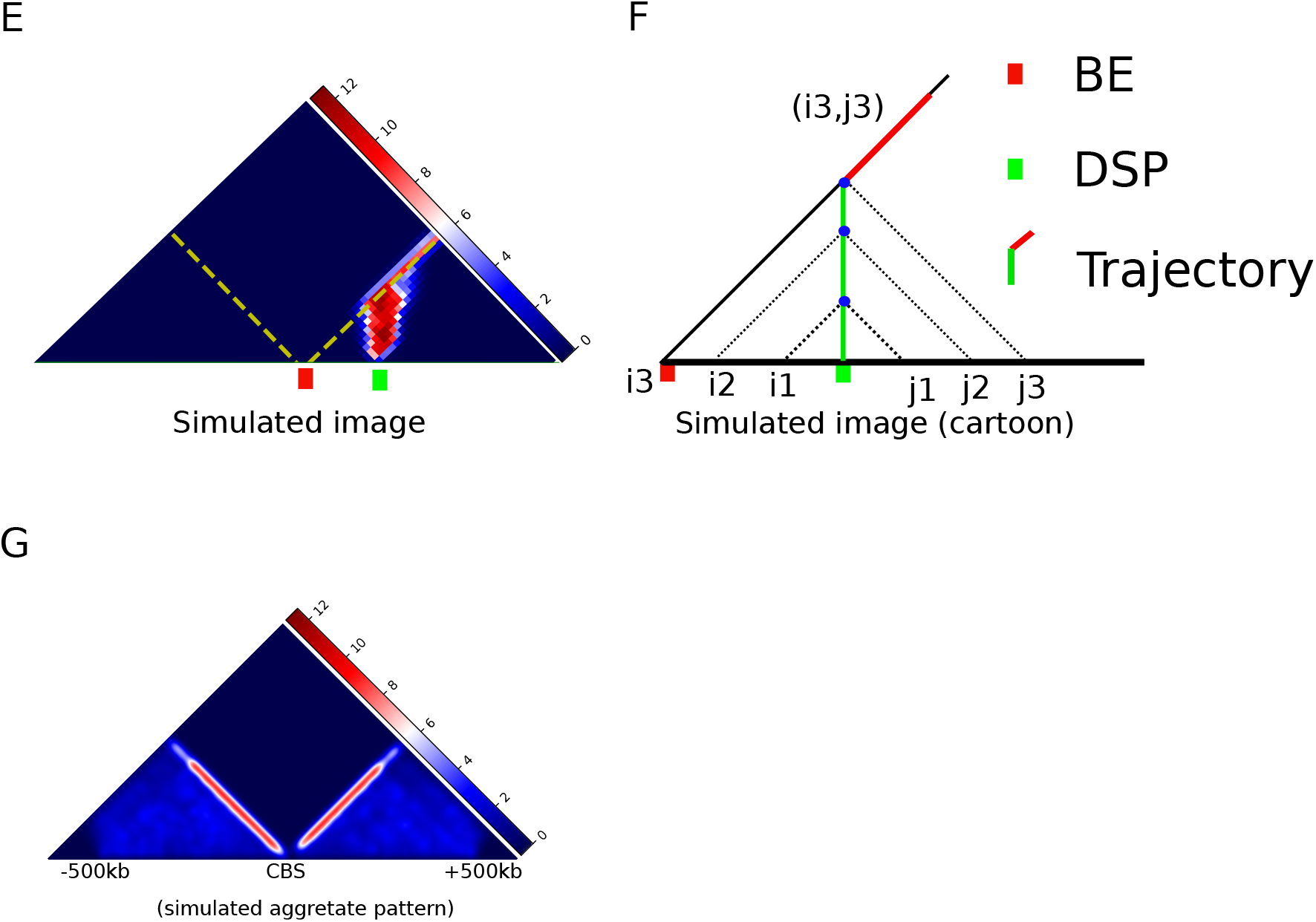
CTCF Aggregation Pattern modeling(CAP). CAP is modeled by aggregating 2000 simulated images generated from the compatible DNA extrusion scenarios. (A) For comparison with modeling, shown is an aggregate Hi-C heatmap derived from published Hi-C data centered on genome-wide CBEs (total 25849 sites) in GM12878 cells. (B) The cartoon shows the composition of the aggregate map, including the SDA, ISD, and SDB. Note that the red line segments represent the two boundaries between SDA/SDB and ISD. (C) The cartoon shows how the domain strip is formed. Top: A loop extrusion event is initiated between a BE (the red rectangle) and several genetic elements (represented by the green, magenta, and blue rectangles). When the BE stops the left anchor, the moving of the right anchor will bring the BE to interact with the downstream genetic elements consecutively. Bottom: the cartoon shows that the DNA extrusion (top panel) is consistent with the domain strip. (D) The flowchart shows the simulation of the aggregate pattern. (E) An example of the simulated Hi-C image corresponding to an ensemble of DNA extrusion trajectories. Note that the DSP (green box) is located at a fixed position at the downstream of the center. (F) The cartoon shows one of the DNA extrusion trajectories in (E). Three dots in the trajectory correspond to three loop configurations whose anchors are located at positions (i1, j1), (i2, j2), and (i3, j3). Note that the red line represents the DNA extrusion motion that the center BE blocks one anchor while the other anchor moves. Most component scenarios share this motion. (G)The heatmap shows the simulated aggregate pattern based on the conditions C1 and C2(See Table 1).

Published data suggest that CAP has two strips that are well consistent with DNA extrusion activities generating the Hi-C feature called a domain strip(Fig2c)[10][24][19]. However, we were not able to recognize the strip features of CAP from a large majority of the component local Hi-C maps(FigS1a). Therefore, two questions needed to be answered to address this discrepancy. First, are the strips in CAP linked to DNA extrusion? Second, can aggregation enhance the features that are missing from the component maps?

We subsequently addressed these two questions using the following algorithm. First, we generated 2000 images, each obtained from the DESs based on a set of assumptions listed in Table 1. Second, we piled up the 2000 images to reproduce the aggregate patterns. Third, we examined to see whether the simulation has reproduced the significant features of the aggregate Hi-C pattern. If the answer was no, then we adjusted the conditions and repeated the steps. If the feature was present, then we analyzed the biological significance of the reconstructed DESs(Fig2d).

**Table 1:** The conditions defining BE and DSP

|  |  |
| --- | --- |
| C1 | A BE is located at the center, which blocks DNA extrusion initiated from the upstream and downstream |
| C2 | A DSP is uniformly located upstream or downstream of the center |
| C2_M | The distribution of DSP is skewed toward upstream (for G1) or downstream (for G2) of the center |
| C3 | A CBEP is uni-directional, It blocks the DNA extrusion from the downstream, while the DNA extrusion from upstream can traverse |
| C4 | A CBEM is uni-directional. It blocks DNA extrusion from the upstream, while the DNA extrusion from downstream can traverse |
| C5 | A CBEM is located upstream of the center CBEP. The position of the CBEM and the center CBEP is not fixed. No DSP is located between this CBEM and the center CBEP. |
| C6 | A CBEP is located downstream of the center CBEM. The position of the CBEP and the center CBEM is not fixed. No DSP is located between this CBEP and the center CBEM. |
| C7 | The center is G1 or G2 promoters. The DNA extrusion is stopped by the neighboring CBEM (for G1) or CBEP (for G2) |
| C8 | A DSP is located at the center, with BEs evenly distributed at two sides |

**Table 2:**
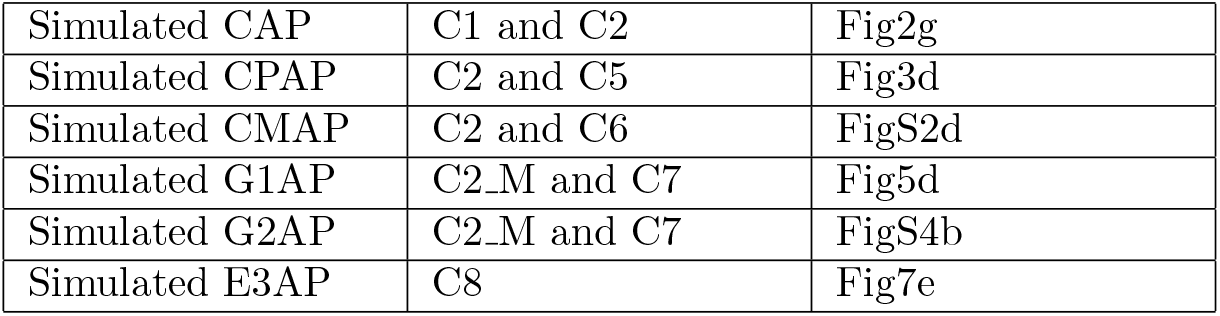
The conditions used for simulating the aggregate patterns

|  |  |  |
| --- | --- | --- |
| Simulated CAP | C1 and C2 | Fig2g |
| Simulated CPAP | C2 and C5 | Fig3d |
| Simulated CMAP | C2 and C6 | FigS2d |
| Simulated G1AP | C2_M and C7 | Fig5d |
| Simulated G2AP | C2_M and C7 | FigS4b |
| Simulated E3AP | C8 | Fig7e |

For illustration, we generated a simulated image based on condition C1 (See Table 1) and an extra condition that a DSP is located at a fixed position downstream of the center (Fig2e). Condition C1 corresponds to the requirement of CAP that a BE (the CTCF binding element) is located at the center. As discussed above, the MC sequence associated with the simulated image collects DNA extrusion trajectories. In this example, all these trajectories began at a fixed DSP. Examining one of these trajectories showed that it was divided into two segments (The red line segment versus the green line segment, in Fig2f). The first segment (Green line segment, Fig2f) represents the DNA extrusion motion within the region of SDB, which is dependent on the scenario-specific DSP. The second segment (red line segment, Fig2f) represents the DNA extrusion motion when the left anchor is blocked by the center CBE while the other anchor continues to move. Thus, the first motion (green line segment) contributes to the signals in SDB (this example), and the second motion contributes to the signals within the boundary to form the strip. Moreover, the center BE prevents the trajectories from entering ISD, which causes the signal level in ISD to be lower than SDB and thus creating a boundary.

The above analysis shows that condition C1 is essential to form the strip feature, with the DSP distribution affecting the signal levels of SDB and SDA. Therefore, we introduced the second condition, C2, which requires that DSPs are uniformly distributed upstream and downstream of the center. Finally, with the combined C1 and C2 conditions, we established DESs which allows the generation of significant features of CAP(Fig2g).

After randomly selecting and analyzing simulated images from the total collection, FigS1b), we observed that the DNA extrusion motion in most scenarios could be divided into two segments akin to the above exemplary image. Aggregation affects signals differently, and thereby forms the strip by highlighting signals that are distributed along the boundary whose pixels are far fewer than the entire SDB. In contrast, all the signals triggered by the DNA extrusion motion within the SDA or SDB are averaged by aggregation, which suggests that these signals have no particular pattern in SDA or SDB. For the same reason, polymer interaction only affects the background signal after aggregation.

Of note, none of the simulated images are derived from an authentic DES based on the real Hi-C data, which is generally associated with a much more complicated trajectory due to the multiple DSPs and BEs. However, in our model, the authentic DES did not cause patterns in SDA or SDB after aggregation, which substantiates our method that is consistent with the aggregation process without modeling the local specific features. Our model, therefore, aims to recapitulate the featured DNA extrusion motion shared by the authentic component scenarios. For example, in CAP, the featured DNA extrusion motion is that one anchor is blocked by the center CBE, while the other anchor continues to move.

Altogether, we confirmed that the two strips of CAP can be the result of DNA extrusion. We also have explained the role of aggregation in enhancing the missing features from the component images, which are relevant to the DNA extrusion motion shared by most component scenarios. Here, we used CAP to demonstrate a method for reconstructing DESs from an aggregate pattern. In the following, we further show that our method can capture more subtle yet essential features of the aggregate patterns indicated by the underlying DESs.

### 3.3 Divergent CBE pairs are necessary for the aggregate patterns centered on CBEs with plus or minus motifs

The CAP discussed above is symmetrical relative to the center CBE. However, previous studies demonstrated that the CBEs with the plus motif (CBEP) or the minus motif (CBEM) interact with other genomic regions with a robust directional bias[11][21]. To investigate the relevant DESs, we generated the CTCF plus aggregate pattern (CPAP, Fig3a) and CTCF minus aggregate pattern (CMAP, FigS2a). CPAP and CMAP show the following features: the same level of signals in SDA and SDB, two boundaries, and only one strip. Of note, CPAP has the strip between SDB and ISD while CMAP resides between SDA and ISD.

**Figure 3:**
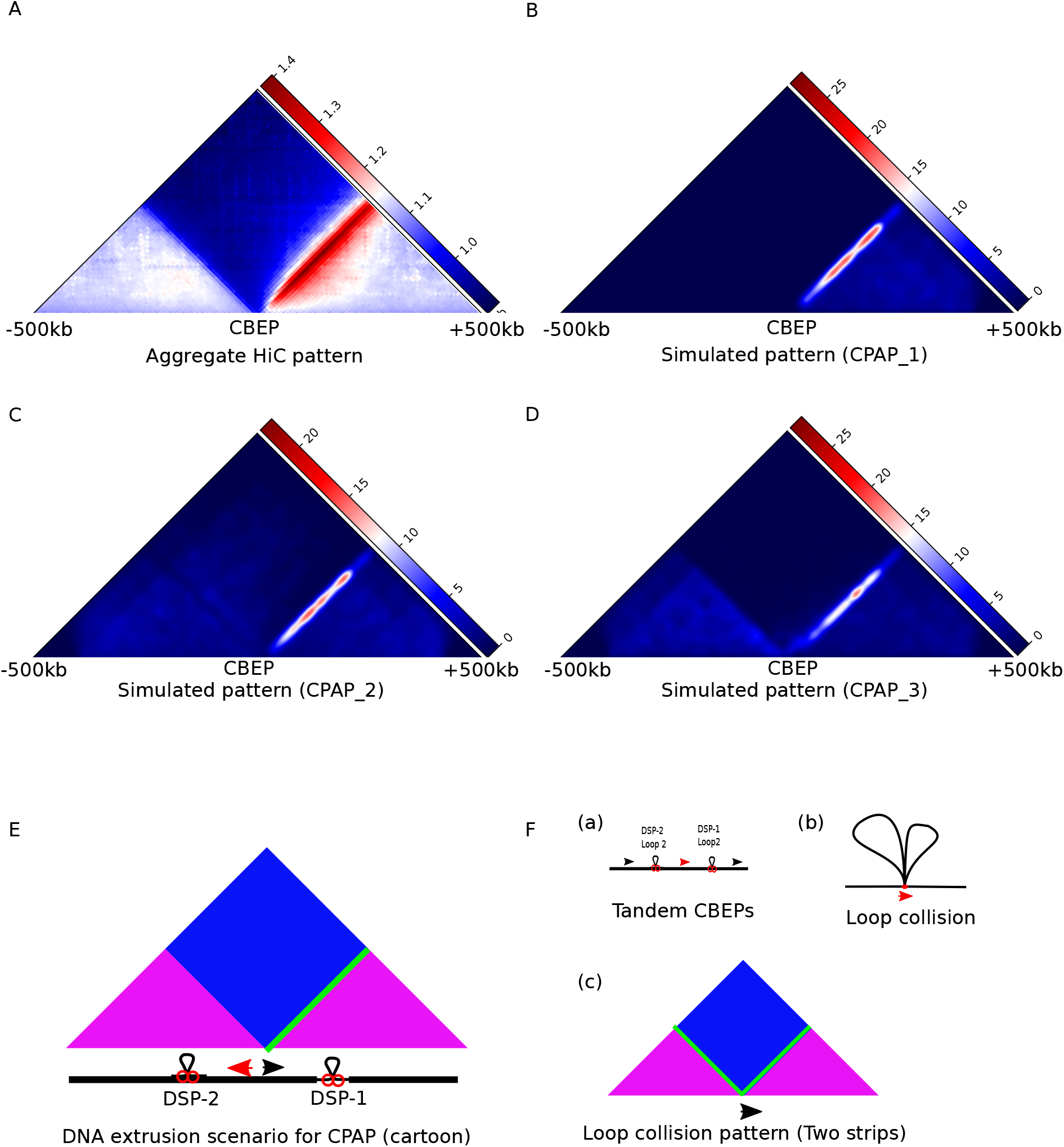
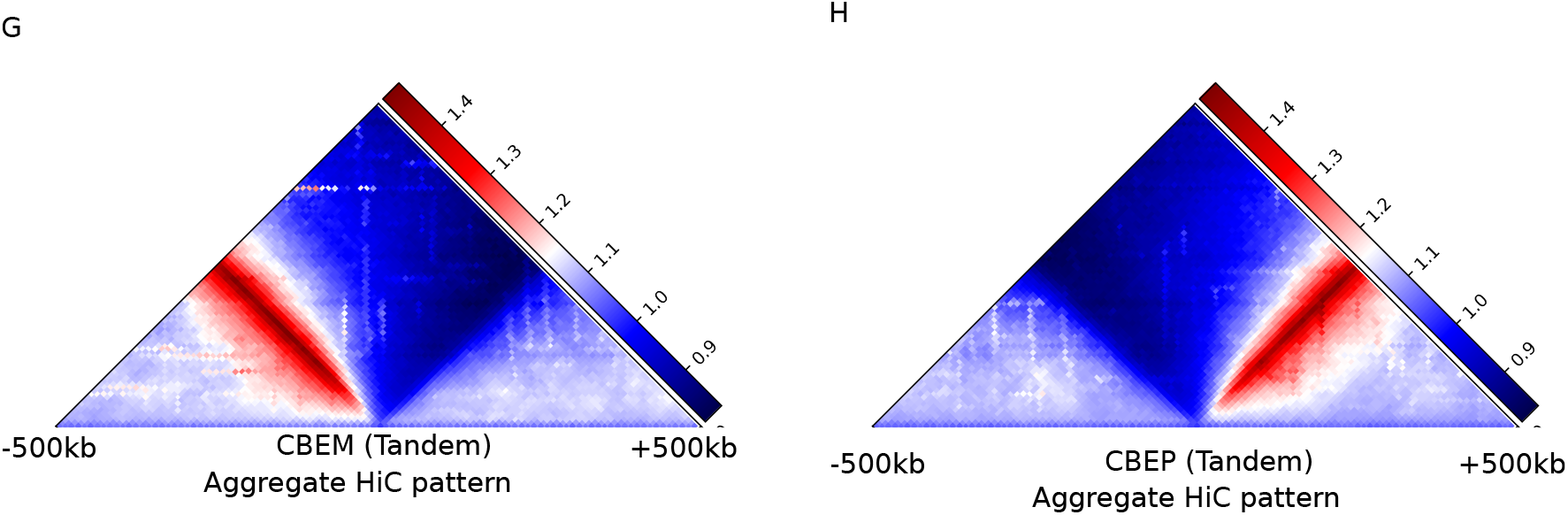
Modeling CPAP. (A)The heatmap shows the aggregate Hi-C map centered on the genome-wide CBEPs. (B) The simulated pattern is generated by the DNA extrusion scenario in which no DNA extrusion events are initiated from upstream of the center. Note that this pattern does not have signals in SDA, which is different from the pattern in (A). (C) The simulated pattern is generated by assuming that the upstream DNA extrusion can traverse the CBEP at the center. Note that it loses the boundary between SDA and ISD. (D) This pattern is generated based on C2 and C5. This pattern reproduces the significant features in (A). (E) The cartoon shows the DNA extrusion scenario for CPAP. Note that the DNA extrusion from the DSP-2 does not access the center as it is blocked by the upstream CBEM (red arrowhead). The DNA extrusion from the downstream DSP-1 forms the strip. (F)The cartoon shows the loop collision model. (a) Two loops are initiated from DSP-1 and DSP2 within the regions containing tandem CBEPs (b) Loop1 and loop2 in (a) may collide at the CBEP (red arrowhead) due to the spatial hindrance. (c) The aggregate pattern consistent with loop collision shows two strips. (G)The aggregate pattern centered on the tandem CBEM in GM12878. (H) The aggregate pattern centered on the tandem CBEP in GM12878. Note that only one strip is present in (G) and (H).

In the following, we used CPAP to illustrate how the underlying DESs are reconstructed. The feature that SDA and SDB have the same level of signals suggests that condition C2 (See Table 1) is still necessary for the scenario. We confirmed this by simulating the DES based on condition C1 and the condition different from C2 that DSP is only located downstream of the center. This scenario generated a pattern showing one strip located at the expected position but without all the signals within SDA (Fig3b). Therefore, we found that condition C1 does not fit the CPAP (and CMAP) scenario, as it generates two strips and cannot be compensated for by adjusting the position of DSP. If we require that DSP is only located at one side of the center, the signals in SDA or SDB will disappear.

We changed condition C1 to condition C3, which requires that the DNA extrusion initiated from the downstream is blocked by the center CBEP, while the DNA extrusion from the upstream can traverse the center CBEP (C3, See Table 1). Condition C3 is consistent with the finding showing that loop traversing is observed in vitro[25]. Based on conditions C2 and C3, we modeled the scenarios which generate a pattern exhibiting the single-strip feature as expected but without the boundary between the ISD and SDA(Fig3c).

Finally, we established a scenario based on condition C2 and a new condition C5(See Table 1). C5 requires that a CBEM be upstream of the center CBEP and no DSP between the upstream CBEM and the center CBEP. Note that C5 is not contradictory with C3. In other words, if C5 is valid, we are unable to judge if C3 is correct since no DNA extrusion initiated from upstream can access the center CBEP. We found that the major features of CPAP can be reproduced by the scenario based on C2 and C5 (Fig3d). In this scenario, the upstream CBEM is not fixed relative to the center, and so the DNA extrusion signals cannot form the strip between SDA and ISD. On the other hand, the upstream CBEM (condition C5) prevents the upstream DNA extrusion from entering the ISD, and so the boundary between SDA and ISD can still be formed. The simulation of CMAP is based on the condition C2 and C6 (see FigS2b to FigS2d, and See Table 1).

The above analysis highlights divergent CBE pairs as the essential structure to generate CPAP and CMAP. In addition, the analysis demonstrates that CBEP and CBEM are only affected by the DNA extrusion activities initiated from one side, confirming the directional bias (Fig3e). Of note, the divergent CBE pair is not consistent with the loop collision model [7][26] because this model is usually associated with two strips (Fig3f). Loop collision was proposed to induce the loop hubs in the regions containing the tandem CBE array (a cluster of CBE with the same motif direction). However, our analysis suggests that the tandem CBE array is associated with one dominant DNA extrusion event rather than multiple; otherwise, the pattern with two strips will emerge. To confirm our analysis, we collected CBEPs and CBEMs that are located in the middle of genome-wide tandem CBE arrays (See the red arrowhead in Fig3f(a)) in GM12878 cells. These CBEPs and CBEMs are about 21% of all CBEs in the whole genome. Consistently, the corresponding aggregate Hi-C patterns show only one strip(Fig3g and Fig3h).

### 3.4 Half of the promoters show strong directional bias to interact with enhancers

We have shown that the reconstructed DESs can explain the directional bias of CBEP and CBEM. Next, we aimed to establish DES association with promoters, which may reveal the formation of promoter enhancer interactions (PEIs). We first generated the promoter aggregate patterns in several different cellular contexts(Fig4a, FigS3a, and FigS3b). The promoter aggregate patterns are similar to the results reported previously [22][23], which have no directional bias.

**Figure 4:**
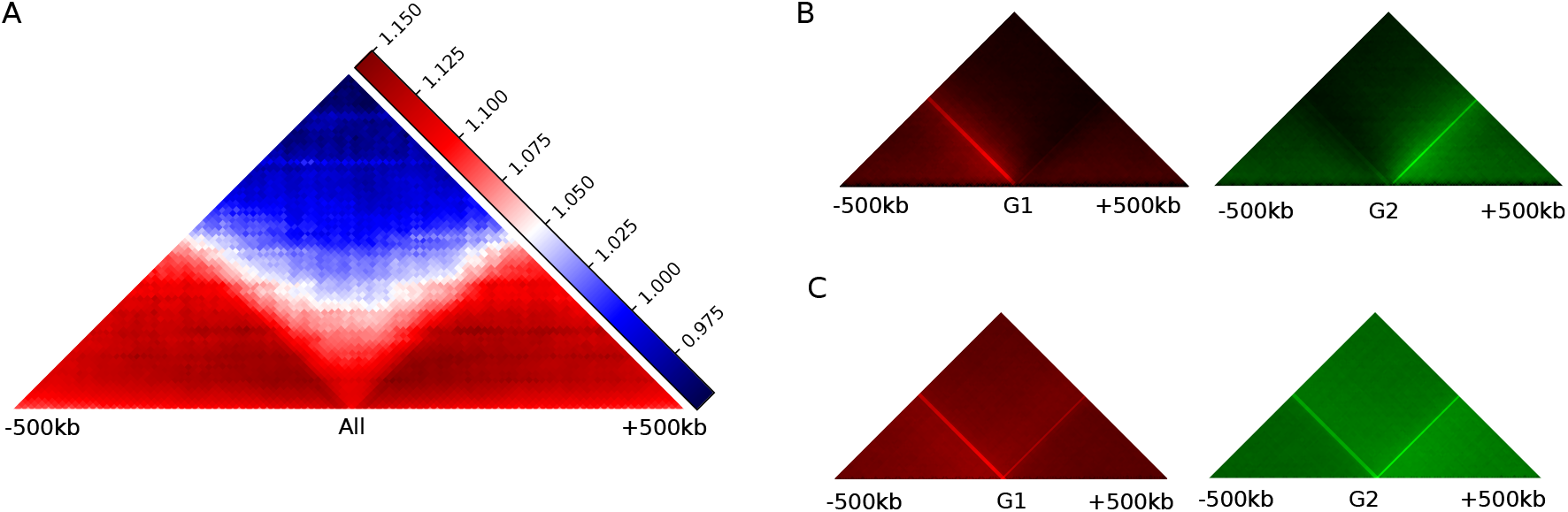
Promoters (G1 and G2) interact with enhancers in a specific direction. (A) The aggregate Hi-C map that is centered on all promoters in GM12878. (B) The directional bias of PEIs for G1 promoters(left, red color) and G2 promoters (right, green color). Note that only the PEI signals are aggregated. For G1 promoters, the left signals are 5.4 fold more than the right signals. (C) The distribution of the enhancer density for G1 promoters (left, red signal) and G2 promoters (right, green signal). For G1 promoters, the left enhancers are 1.4 times more than the right enhancers.

Since CPAP and CMAP show a directional bias while the CAP combining CPAP and CMAP does not, directional bias may only be associated with a subset of promoters. To determine a promoter’s directional bias, we computed a value (Directional Bias index, DBI) which is equal to the difference between the number of upstream promoter-enhancer interactions (PEIs) and the downstream PEIs (See methods for the full definition of PEIs). Promoters that only contacted one enhancer are excluded from analysis, since this interaction alone does not assess their directional bias. Based on the DBI, the entire promoters could be categorized into three groups - G1 (16715 counts), G2(17598 counts), and G3 (35833 counts). The G1 promoters have the positive DBI, and the G2 promoters have the negative DBI. G3 consists of all other promoters that are not G1 or G2 (See supplementary list PromoterGroup).

Consistent with the DBI, the PEI aggregate signals centered on the G1 and G2 promoters show strong directional bias (Fig4b). The directional bias is not due to the imbalanced distribution of enhancers relative to the promoters. We have observed 5.4 times more signals on the stronger side than the weaker side for G1 promoters(Fig4b, left panel), whereas there are only 1.4 fold more enhancers located upstream than the downstream for the G1 promoters (Fig4c, left panel, p *<* 1e-16, binomial distribution). A similar finding was observed for G2 promoters (Right panel in Fig4b and Fig4c). Additionally, G1 and G2 promoters have significantly higher expression levels than the G3 promoters indicating that the directional bias is important for the promoter’s activity (FigS3c).

### 3.5 The directional bias of G1/G2 promoters is dependent on the neighboring CBEs

In addition to the aggregate PEI pattern (Fig4b), we next generated the aggregate Hi-C patterns centered on G1 promoters (G1AP, Fig5a) and G2 promoters (G2AP, FigS4a). We found that the overall structure between G1AP and CMAP (FigS2a) is quite similar, indicating that these two patterns have strong connections. Similar relationships can be observed between G2AP and CPAP. However, G1AP and G2AP show a more diffusive strip than CMAP and CPAP, suggesting the underlying DESs are not equivalent.

**Figure 5:**
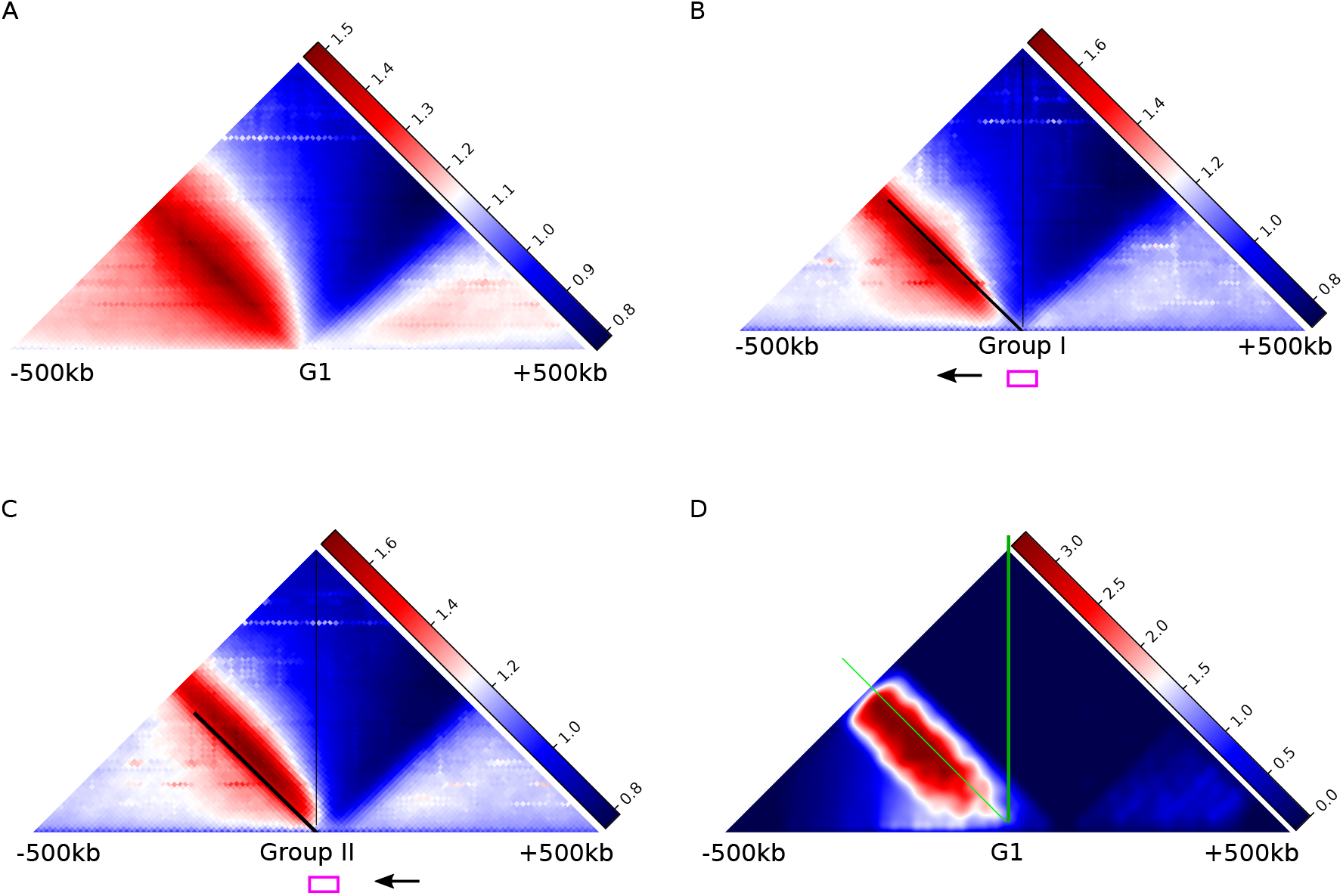
G1AP is related to the neighboring CBEM. (A) The aggregate Hi-C pattern centered on G1 promoters (G1AP). Note that G1AP has a strip more diffusive than that in CMAP (FigS2a). (B) Aggregate Hi-C map centered on Group I G1 promoters. The Group I G1 promoters (represented by the magnet box) locate downstream of the CBEM (represented by the black arrow). Note that the signals are enriched within the SDA, which is left of the boundaries (The black line)(C) Aggregate Hi-C map centered on the Group II G1 promoters. The Group II G1 promoters (represented by the magenta box) locate upstream of the CBEM (represented by the black arrow). Note that the signals are enriched within the ISD, which is right of the boundaries (Black line. (D) The heatmap shows the simulated result for G1AP. Note that the diffusive strip is formed by the neighboring CBEM, not by the center G1 promoters

The center element of the aggregate pattern is essential for reconstructing the associated DES. We first examined the colocalization between G1 promoters and CBEM, the center of G1AP and CMAP, respectively. About 10% of G1 promoters are located at the same genomic bin (10kb) with a CBEM. This number increases to 60% if the colocalization region is expanded to 50 kb. The G1 promoters were divided into two groups based on their positions relative to the closest CBEM. Specifically, group I promoters are located downstream of the CBEM, while the group II promoters are located upstream of the CBEM. The corresponding aggregate patterns show the enriched signals at different sides of the boundary between SDA and ISD (Fig5b and Fig5c). Pattern I (for group I G1 promoters, Fig5b) shows more signals on the left side of the boundary (within the SDA), whereas pattern II (Fig5c) shows more signals on the right side of the boundary (within the ISD). In other words, we found that the enriched signals are more related to the neighboring CBEM rather than the center G1 promoters. The neighboring CBEM may serve as the authentic BE to stop the DNA extrusion, whereas the G1 promoters at the center do not. We finally reproduced the G1AP and G2AP with the corresponding DESs based on conditions C2 M and C7 (Fig5d and FigS4b. See below for more details about using C2 M instead of C2.). Based on the corresponding DESs, the directional bias observed for the G1 and G2 promoters is confirmed by the neighboring CBEM and CBEP, respectively.

### 3.6 The directional bias of G1AP and G2AP is determined by imbalanced DNA extrusion activities

In addition to adjoining CBEMs, G1 promoters also adjoin CBEPs (Fig6a and FigS5b). About 60% of G1 promoters border with CBEM and 33% border with CBEP. A similar pattern is seen for G2 promoters (FigS5a and FigS5c). Here, the neighboring region is defined as the 50kb region flanking the promoters (Same for the below). The DNA extrusion associated with the neighboring CBEPs draws G1 promoters to contact the downstream loci, but the G1 promoters show a strong directional bias to interact with the upstream loci.

**Figure 6:**
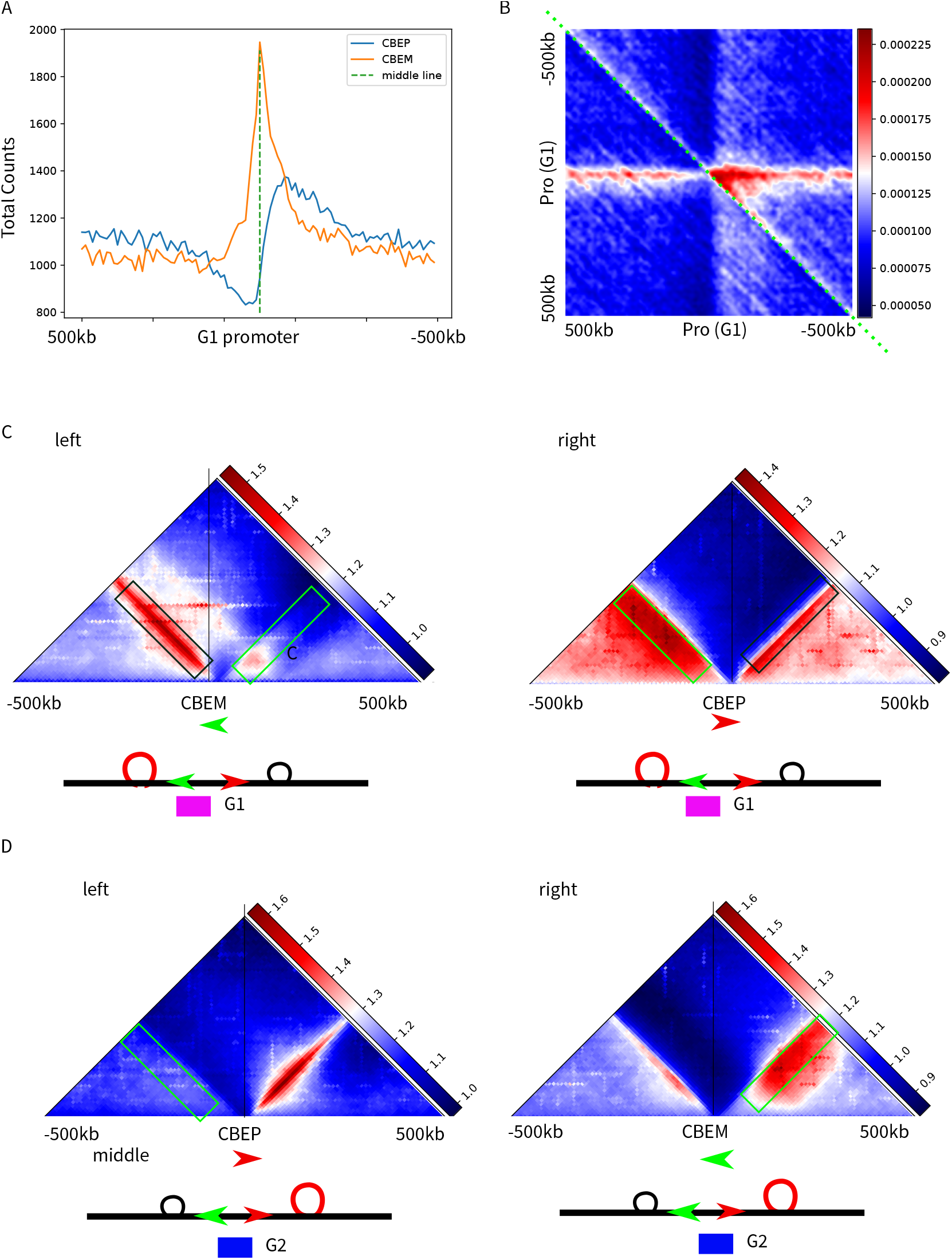
G1 promoters are pulled toward upstream by more vigorous upstream DNA extrusion activities. (A) The line plot figure shows the distribution of CBEM and CBEP relative to the G1 promoters. Note that the CBEM (yellow line) is about two-fold more than the CBEP (blue line). (B) The joint probability heatmap shows the distribution of the divergent or convergent CBE. The enriched high-intensity dots (red signal) locate the downstream of G1 promoters (Center) and represent the divergent configuration (Locate at the upper triangle). (C) The upstream DNA extrusion is more active than the downstream one. The upstream DNA extrusion activities are represented by the signals in the green box in the right panel. The downstream DNA extrusion activities are represented by the signals in the green box in the left panel. (D) Same as (C) but for G2 promoters. The downstream DNA extrusion activities are more potent than those upstream. The red and green arrowheads represent CBE with plus and minus motifs, respectively. The G1 promoters and G2 promoters are represented by the magenta and blue box, respectively.

To investigate this discrepancy further, we first examined the CBEP and CBEM close to G1 promoters to see whether they can form convergent or divergent configurations. The distributions of these two configurations relative to G1 promoters are shown in the joint probability heatmaps (Fig6b, See FigS5d for G2 promoters). For a dot with the coordination i (x-axis) and j (y-axis) in the heatmap, the color intensity indicates the probability of finding the CBEP at the *i*_*th*_ position and a CBEM at the *j*_*th*_ position. Thus, the lower triangle of the heatmap (*j > i*, regions left to the green dash line in Fig6b) represents the probability of the convergent CBE pairs, and the upper triangle (*i > j*) represents the probability of the divergent CBE pairs. According to the distribution of high-intensity dots, we found that the neighboring CBEPs and CBEMs mainly form divergent pairs and are located downstream of the G1 promoters. For G2 promoters, CBEP and CBEM also form the divergent configuration but are located at the upstream(FigS5d).

Both G1 and G2 promoters surround a divergent CBE pair, suggesting that these promoters were affected by DNA extrusion initiated upstream and downstream. So why do G1 and G2 show robust directional bias? As discussed above, the signal level in SDA or SDB is indicative of the DNA extrusion strength upstream or downstream, respectively. We found the signal level in G1AP’s SDA is higher than SDB (Fig5a), indicating that these promoters are affected by two imbalanced DNA extrusion activities, with the upstream being more potent than the downstream.

To further confirm the differing strengths of the upstream and downstream DNA extrusion activities, we generated aggregate patterns centered on the CBEP and CBEM that surround G1 and G2 promoters. As multiple CBEP and CBEM adjoin G1 or G2, we selected the targets for aggregation with the two approaches(FigS5e). First, we chose a pair of divergent CBEs whose linear distance is the largest. Second, we chose all the CBEP or CBEM within the regions (300kb) flanking the center. Of note, the signals within the SDB in the aggregate pattern centered on the CBEM reflects the downstream DNA extrusion activities (Green box in the left panel in Fig6c), while the signals within the SDA in the aggregate pattern centered on the CBEP(Green box in the right panel in Fig6c) represents the upstream DNA extrusion activities. We consistently observed that the upstream DNA activities are more active than the downstream activities for G1 promoters (method1: Fig6c, method2: FigS5f). The converse is true for G2 promoters where the downstream DNA extrusion activities are more active than the upstream ones.(Fig6d)

Taken together, we have demonstrated that the directional bias of G1 and G2 promoters results from the imbalanced DNA extrusion activities associated with the local divergent CBE pairs. While the exact mechanism underlying the imbalanced DNA extrusion activities is not known, our analysis suggests that regulation of the distribution of DSPs may play an important role.

### 3.7 DNA extrusion does not stop at enhancers

Recent studies suggest that DNA extrusion may stop or pause at enhancers and promoters to facilitate promoter-enhancer interactions[23][27]. However, our analysis indicates that one anchor of the DNA extrusion stops at the neighboring CBEs, but not the G1 or G2 promoters. Whether the other anchor stops at the enhancers was further examined.

We analyzed the aggregate maps centered on enhancers by collecting the enhancers that interact with G1 or G2 promoters at a high intensity (*OE >* 1.5, see methods). These enhancers were divided into three categories - E1, E2, and E3 (See supplementary list EnhancerList). E1 enhancers (4027 counts) only interacted with G1 promoters, E2 enhancers (4078 counts) only interacted with G2 promoters, and E3 enhancers (13427 counts) contacted with both G1 and G2 promoters. The aggregate patterns centered on E1, E2, and E3 are named E1AP (Fig7a), E2AP(FigS6a), E3AP (Fig7b), respectively.

**Figure 7:**
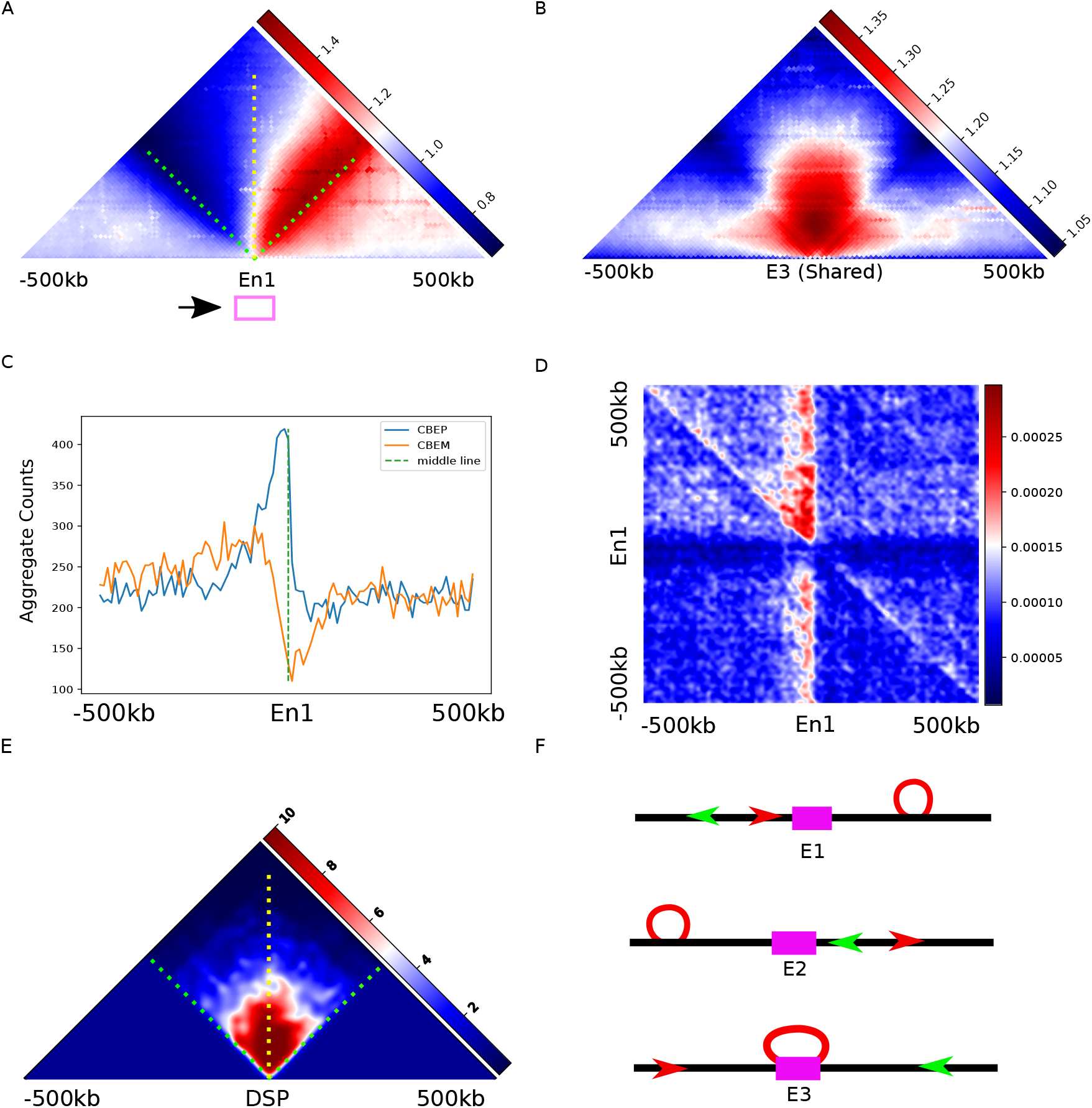
DNA extrusion does not stop at enhancers. (A) The Hi-C heatmap from GM12878 data shows E1AP, an aggregate pattern centered on the E1 enhancers (E1, contact with G1 promoters only). Note that E1AP and G2AP (FigS4a) have similar structure. (B) The Hi-C heatmap shows E3AP, an aggregate pattern centered on the E3 enhancers (the enhancers contact both G1 and G2 promoters). (C) The line plot shows the aggregate CBE signals centered on E1 enhancers. Note that CBEP is distributed more at the upstream side than the downstream side of the center E1 (refer to Fig6a and FigS5a). (D) The joint probability heatmap shows E1 surrounding the divergent CBE pairs (See Fig6b and FigS5d for more details). (E) The simulated aggregate pattern centered on the DNA extrusion starting points (DSP). The overall structure is similar to E3AP. (F) The cartoon shows the DESs associated with E1, E2, and E3 enhancers. The CBEP is represented by the red arrowhead. The CBEM is represented by the green arrowhead. The loop extrusion (red circle) is blocked by the CBE clustering toward E1 (red arrowhead) and E2 (green arrowhead). The E3 enhancers are overlapped with the DSP rather than the BE.

E1AP shows the features similar to G2AP, except for the signals that are more skewed to the ISD side of the boundary. Moreover, the aggregate CBE signals (Compare Fig7c with FigS5a), and the joint probability heatmap (compare Fig7d with FigS5d) are all similar to those of the G2 promoters. A close relationship could also be observed between E2AP and G1AP. Therefore, we conclude that DNA extrusion does not stop at E1 or E2 enhancers but at the neighboring CBEs (Also see FigS6b and FigS6c).

As opposed to E1AP and E2AP, E3AP shows a pattern with the signals enriched within ISD, which is different from all other aggregate patterns. Furthermore, the distribution of CTCF centered on E3 enhancers (FigS6d) is dramatically different from those centered on E1 and E2 enhancers (Fig7c and FigS6b). Therefore, the DES associated with D3 is different from those associated with E2 and E3. Based on the defined transitions between loop configurations (Fig1b), we introduced condition C8, which requires that the center be a DSP (Table 1). The corresponding scenarios based on C8 generated a pattern with the signals enriched within ISD (Fig7e). Altogether, our analysis suggests that E3 enhancers are more likely to be DSPs rather than BEs.

The above analysis indicates that E1, E2, and E3 enhancers do not block DNA extrusion (Fig7f). Therefore, the observed PEIs may be the secondary effect of the convergent CTCF loops formed by DNA extrusion. If this holds true, we should be able to find enhancers that surround one anchor of the CTCF loop while promoters cluster around the other anchor. For G1 promoters, we identified the closest CBEM and the closest upstream CBEP and examined how the enhancers are distributed within this convergent CTCF domain (FigS6e). For three defined regions (50kb) that flank the upstream CBEP (Region 1), the middle position (Region 2), and the downstream CBEM, we found enhancers that are more enriched within regions 1 and 3 but relatively depleted within region 2(FigS6f). A similar finding was observed for G2 promoters(FigS6g). Therefore, the distribution of enhancers supports the hypothesis that PEIs rely on convergent CTCF loops.

## 4 Discussion

This study has introduced a new method for reconstruction of DESs from aggregation patterns. Our analysis demonstrated that aggregation enhances the signals generated by genome-wide, commonly shared DNA extrusion movements, whereas it levels out those caused by specific local DNA interactions. Furthermore, our modeling has successfully reproduced several aggregate patterns suggesting that the underlying assumptions are reasonable, and this method can recapitulate the aggregation effect.

Besides the DNA extrusion model, PEIs could be explained by the polymer interaction or random diffusion model. However, PEIs formed via DNA extrusion are likely to be more stable and efficient as the biological information coded in one-dimensional DNA sequences could direct three-dimensional chromatin interactions[7]. Currently, two assumptions exist to explain the formations of PEIs by DNA extrusion. The first is that DNA extrusion pauses or stops at the promoter and enhancers, which generates loops anchoring on the promoter and enhancer directly[27]. The second assumption is that DNA extrusion forms the convergent CTCF loops, bringing the promoters and enhancers close but not in contact directly[28]. Resolving the discrepancy between these two assumptions is not a trivial task.

Our data support the latter view because the corresponding DESs in our study suggest that DNA extrusion does not stop at promoters or enhancers, but stops only at the neighboring CBEs. Of note, our findings cannot exclude the possibility that promoters or enhancers serve as weaker stop signals. If this is true, our method cannot reveal the patterns formed by promoters and enhancers as the patterns formed by the neighboring CBEs are too strong. Unfortunately, pursuing a Hi-C map with a higher resolution cannot resolve this issue as the “overwhelming effect” still exists[27][23]. In contrast, the aggregate map based on the Hi-C data with CTCF depleted may provide additional information.

Interestingly, we demonstrated that some enhancers (E3 enhancers) contacting G1 and G2 promoters may serve as DSPs. Therefore, a multiple loop structure that encloses several enhancers and promoters can be formed. Of note, this loop structure does not anchor on promoters or enhancers.

Our study highlighted that the DSP plays an essential role within a specific DES. The mechanism which regulates the DSP at both sides of G1 and G2 promoters explains why the imbalanced DNA extrusion activities exist. Evidence points to the loading factor Nipbl, the local transcription, and the releasing factor Wapl as mediators for DSP[14][15]. DSP may also contain the consensus sequences, which are important targets for future genome editing studies. It will be interesting to know whether regulating DSP distribution could affect PEIs, DNA recombination, and other genome folding events.

A recent study demonstrated that loops colliding at a CBE can form enhancer hubs that bring promoters and multiple enhancers together[26]. However, our analysis indicated that this scenario does not happen frequently as it will generate two strips in CPAP and CMAP. For the same reason, we propose that the tandem CTCF plus or minus binding sites are associated with a dominant DNA extrusion event but not with multiple ones. Our postulation is consistent with the DES reported for the protocadherin alpha gene. In this gene locus, only one promoter can interact with the distal enhancer region, suggesting that a dominant DNA extrusion event is responsible for the PEI and no hub structure is formed[29]. This scenario may be valid for other genomic loci containing the tandem CBEs, such as the IGH region in which the V(D)J recombination is implemented[30][31].

Finally, it is essential to note that our conclusion is based on the aggregate patterns that reflect the average behaviors of many DNA extrusion events. Thus, we anticipate more upcoming studies about the specific DES that underlies the local PEIs or other vital processes.

## 5 Methods

### 5.1 Hi-C data processing and constructing the aggregate map

Hi-C data were obtained from GEO as shown in table 3 and processed according to the method described in [3]. First, the sequencing data were aligned to hg19 or mm10, and the low mapping quality reads were removed (MAPQ *<* 30). Second, the data were filtered for the duplicates, the same fragment reads, and the dangling ends. Finally, the raw matrices were represented at the 10kb resolution and normalized by the method described in[32]. The Observed/Expected matrices were obtained using the method described in [1]. The aggregate maps were generated on the O/E matrices using the method described in [3].

**Table 3:**
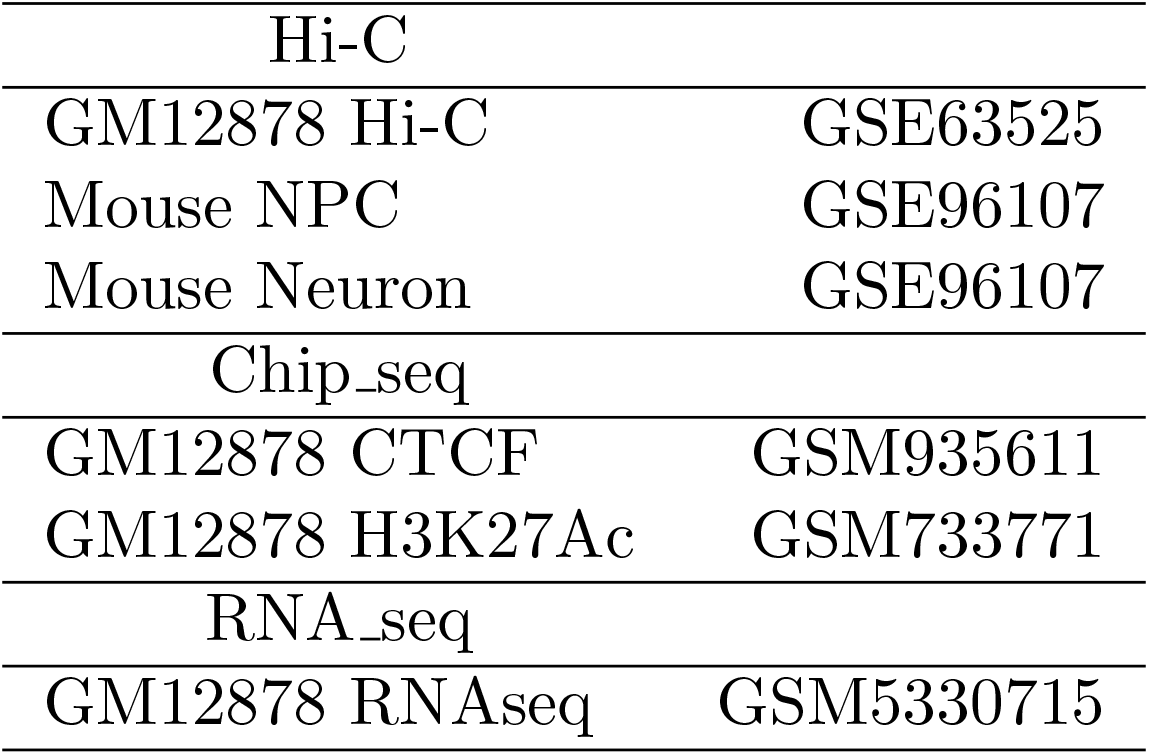
Datasets

| Table 3: Datasets |  |
| --- | --- |
| Hi-C |  |
| GM12878 Hi-C | GSE63525 |
| Mouse NPC | GSE96107 |
| Mouse Neuron | GSE96107 |
| Chip-seq |  |
| GM12878 CTCF | GSM935611 |
| GM12878 H3K27Ac | GSM733771 |
| RNA-seq |  |
| GM12878 RNAseq | GSM5330715 |

### 5.2 Modeling DNA extrusion by Markov Chain

We modeled DNA extrusion as the following. First, the DNA undergoing DNA extrusion is defined as a one-dimensional lattice structure composed of 100 units, and each indexed unit represents a 10kb genomic bin. Second, as the anchors of the loop move along the DNA lattice during the DNA extrusion, their absolute position, represented by a pair *S*_*i,j*_ = (*i, j*), constitutes a state of a stochastic model, and corresponds to the loop configuration in which the left and right anchors are located at positions *i* and *j* respectively, where 0 *<*= *i, j <*= 99 and *i < j* for all *i* and *j*.

Assuming the left and the right anchors do not shift, the induced state space contains 4950 individual states because for every fixed *i*, there are 100 − (*i* + 1) unique values of *j*.

We then defined a discrete-time Markov Chain over a sequence of transition events {*E*_*t*_}, with *t* ∈ ℤ, given by the following transition scheme if no BE (insulator) is encountered.

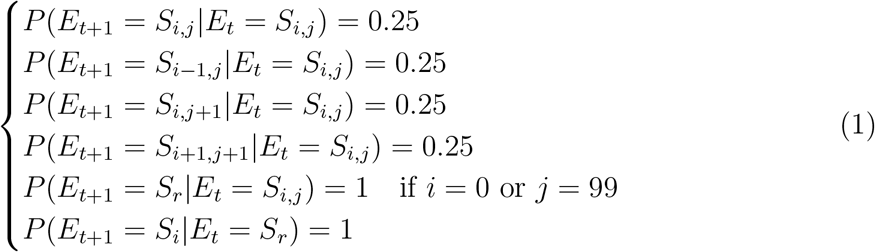

If BE is encountered

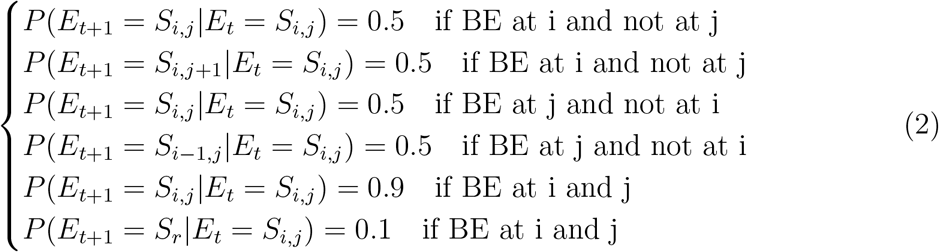

where *E*_0_ = *S*_*i*_ corresponds with LEF binding to DNA, and the two anchors are located at two adjacent positions, and *S*_*r*_ represents the configuration where LEF is released from DNA.

### 5.3 CBEP and CBEM

The CTCF chip seq dataset is obtained from GEO (table 3). HOMER is used to identify the plus or minus motif in a peak region. The list of CTCF binding sites with plus or minus motif is shown in the supplementary list CtcfMotifDirection.

### 5.4 Definition of PEIs

PEIs are the Hi-C interactions between a promoter region and an enhancer region, and the O/E strength (Observed/Expected) is above 1.5. A promoter region is the 10kb genomic bin containing the transcriptional starting sites of the annotated promoters. An enhancer region is the 10kb genomic bin containing the middle point of the peaks of the histone marker H3K27Ac (table 3).

## 7.1 Abbreviation

CTCF: CCCTC-binding factor
TAD: topological associated domain
LEF: loop extrusion factors
BE: barrier element
CBE: CTCF binding sites
CBEM: CTCF binding sites with minus motif
CBEP: CTCF binding sites with plus motif
CAP: CTCF aggregate pattern
CPAP: CTCF plus aggregate pattern
CMAP: CTCF minus aggregate pattern
G1AP: G1 promoter aggregate pattern
G2AP: G2 promoter aggregate pattern
PEI: promoter enhancer interaction
E1AP: E1 aggregate pattern
E2AP: E2 aggregate pattern

## 7 Appendix

### 7.2 Supplementary Figures

**Figure S1:**
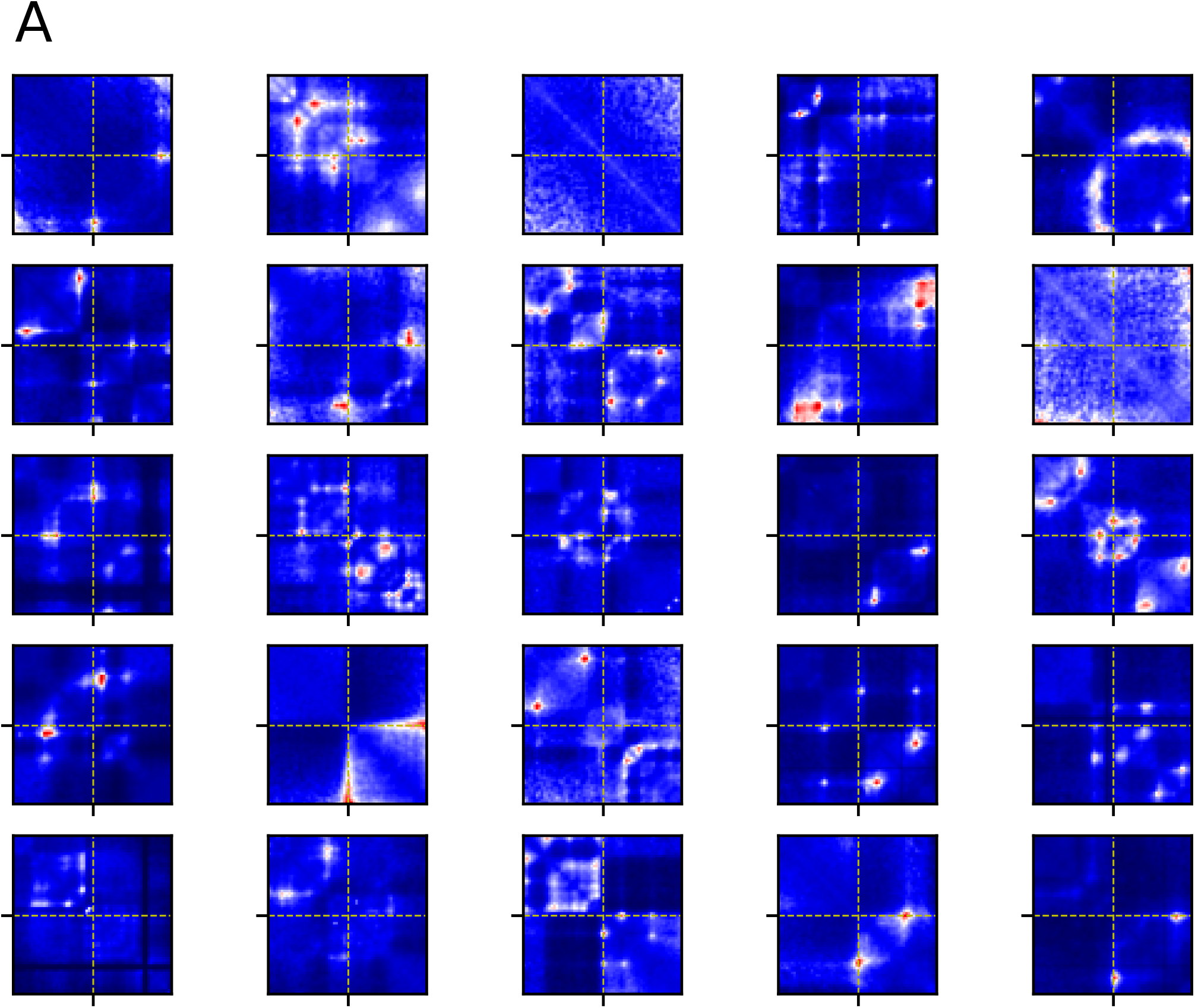
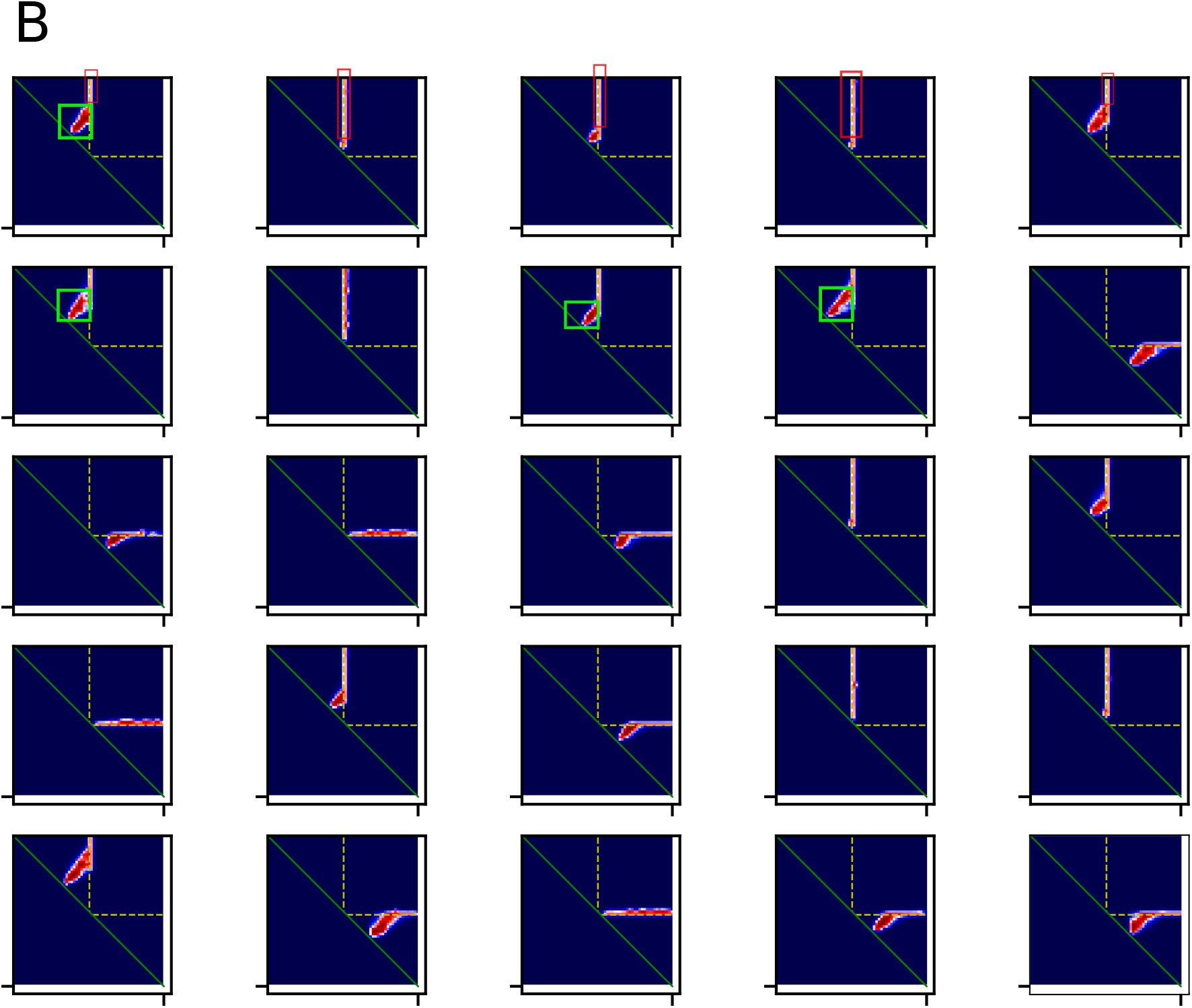
(A) The randomly selected component images which have been aggregated to generate CAP(Fig2a). Note that these images do not show the features in CAP. (B) The images were randomly selected from the collection of 2000 simulated images aggregated to reproduce the CAP. Note that most of the images have two types of signals. The first type, highlighted by the green box, is specific to the local region and averaged during aggregation. The second type, highlighted by the red box, is kept and enhanced to form the strip. On the other hand, the center CBEs prevent the DNA extrusion from moving into the ISD (Between two yellow dash lines). Therefore, the boundaries between the SDA/SDB and ISD are formed.

**Figure S2:**
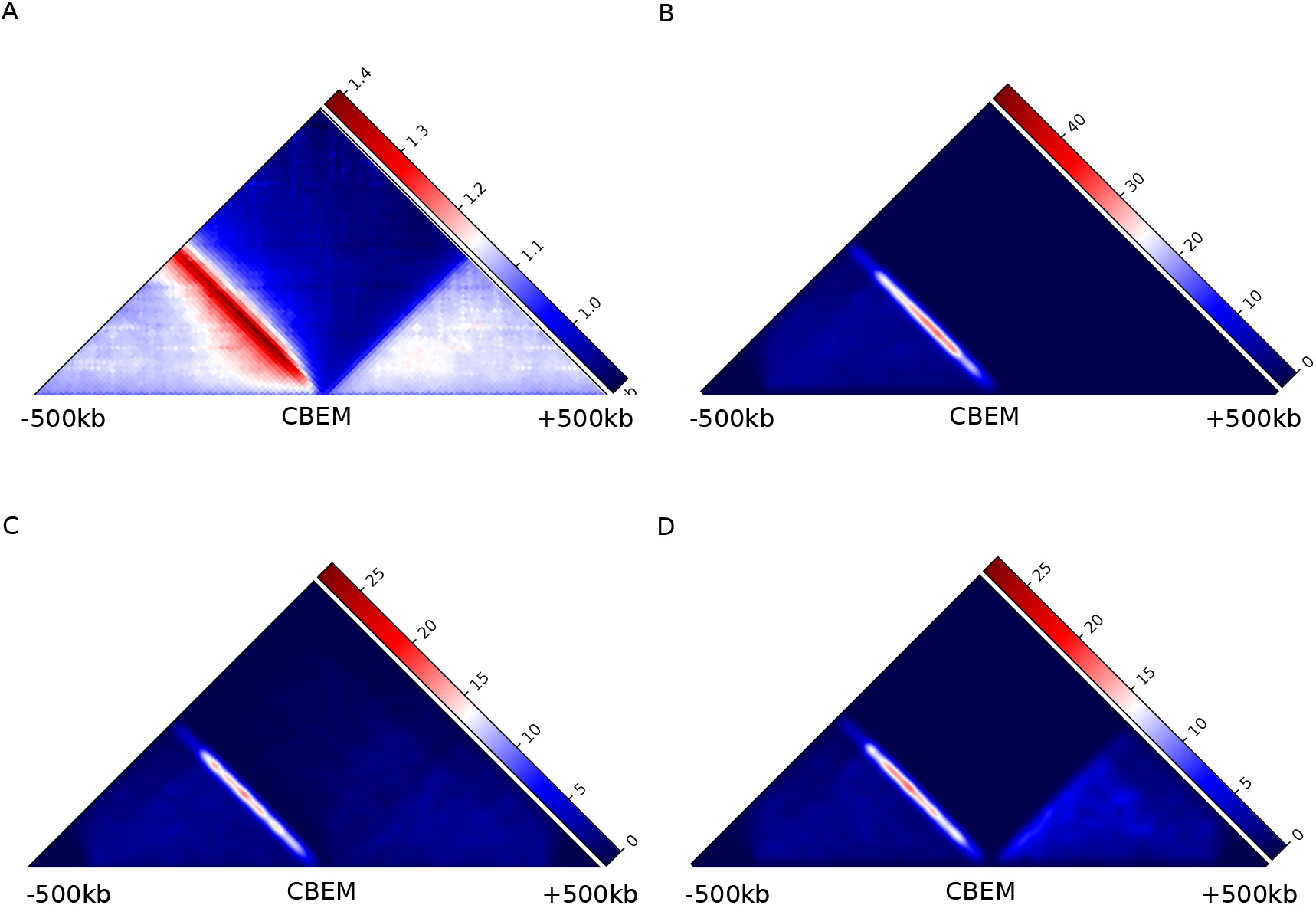
Modeling the CMAP. Refer to the main figures from Fig3a to Fig3d. (A) The aggregate map centered on the genome-wide CBEMs. Note that this pattern has a strip between SDA and ISD, which is different from CPAP (Fig3a). (B) The simulated pattern is generated by the DNA extrusion scenario in which no DNA extrusion events are initiated from the downstream of the center. Note that this pattern does not have signals in SDB. (C) The simulated pattern is generated by assuming that the DNA extrusion initiated from the downstream can traverse the CBEM at the center. Note that it loses the boundary between SDB and ISD. (D) This pattern shows the major features similar to CMAP.

**Figure S3:**
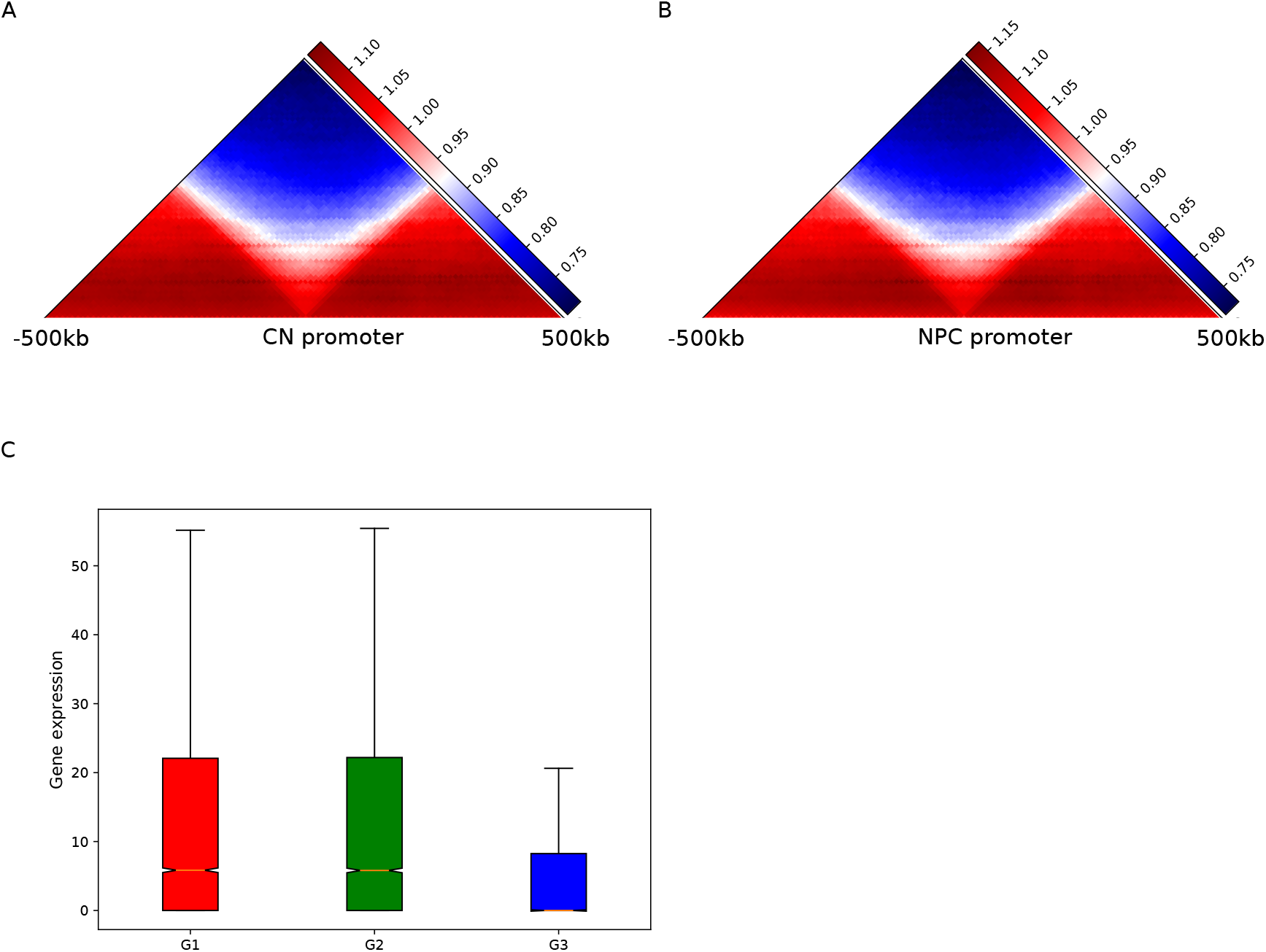
(A) Aggregate Hi-C map centered on promoters in mouse central neuron cells. (B) Aggregate Hi-C map centered on promoters in mouse neural progenitor cells. (C) The box plot figure shows the gene expression level associated with G1, G2, and G3 promoters in GM12878 cells.

**Figure S4:**
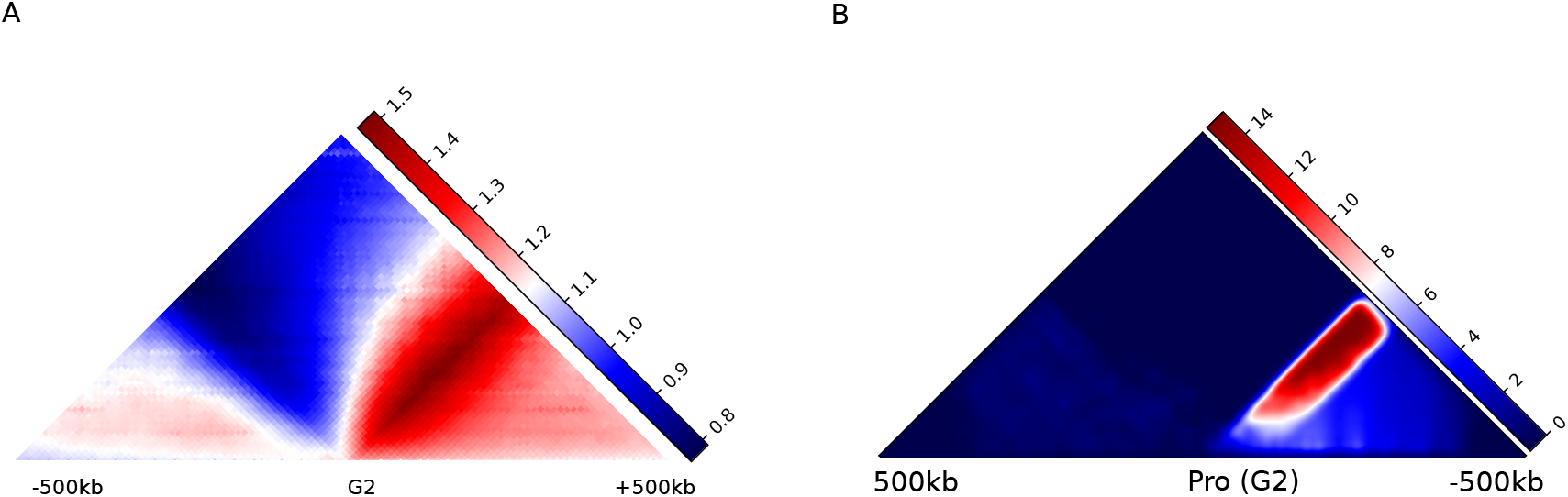
(A) The aggregate Hi-C pattern centered on G2 promoters. (B) The simulated G2AP.

**Figure S5:**
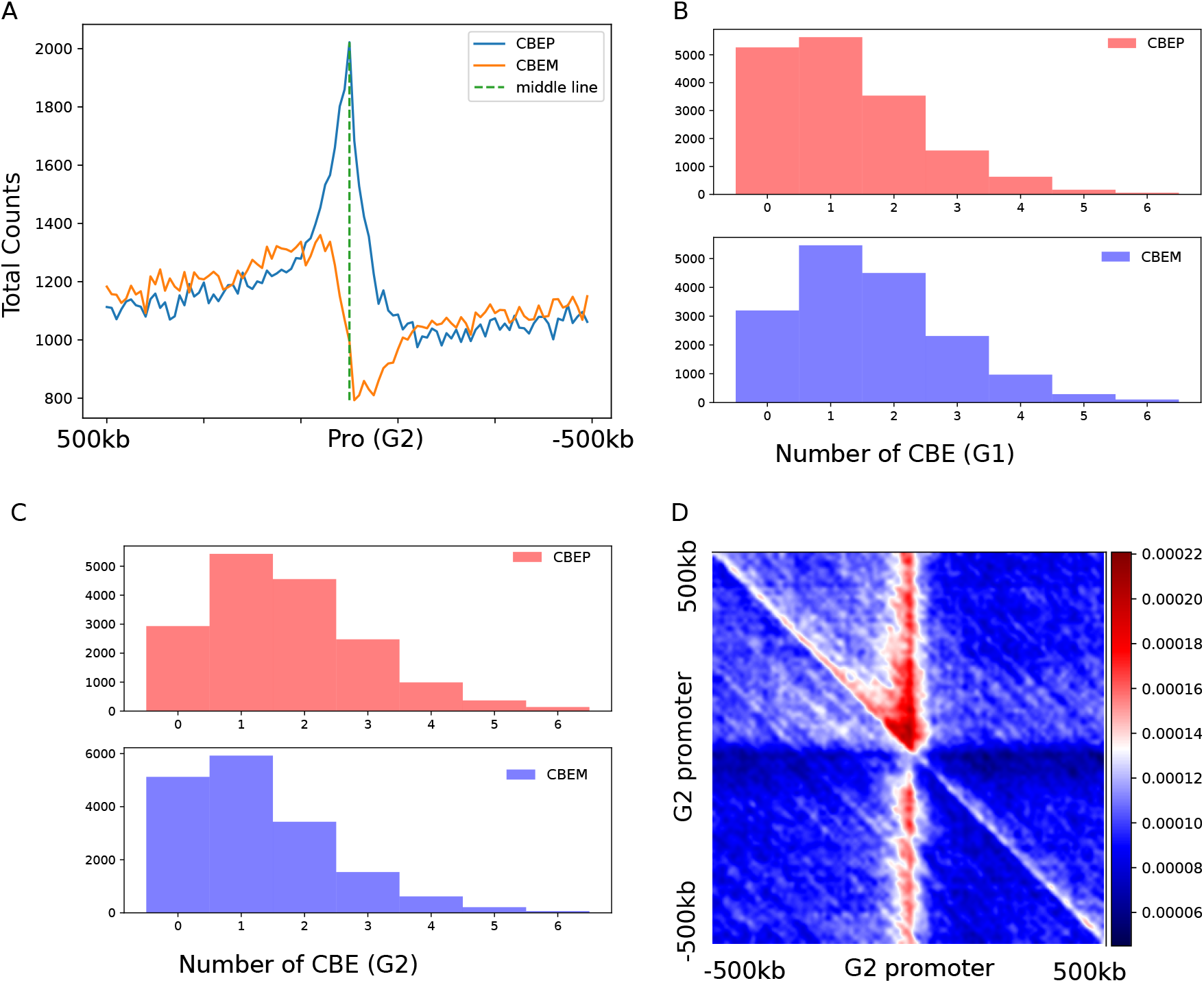
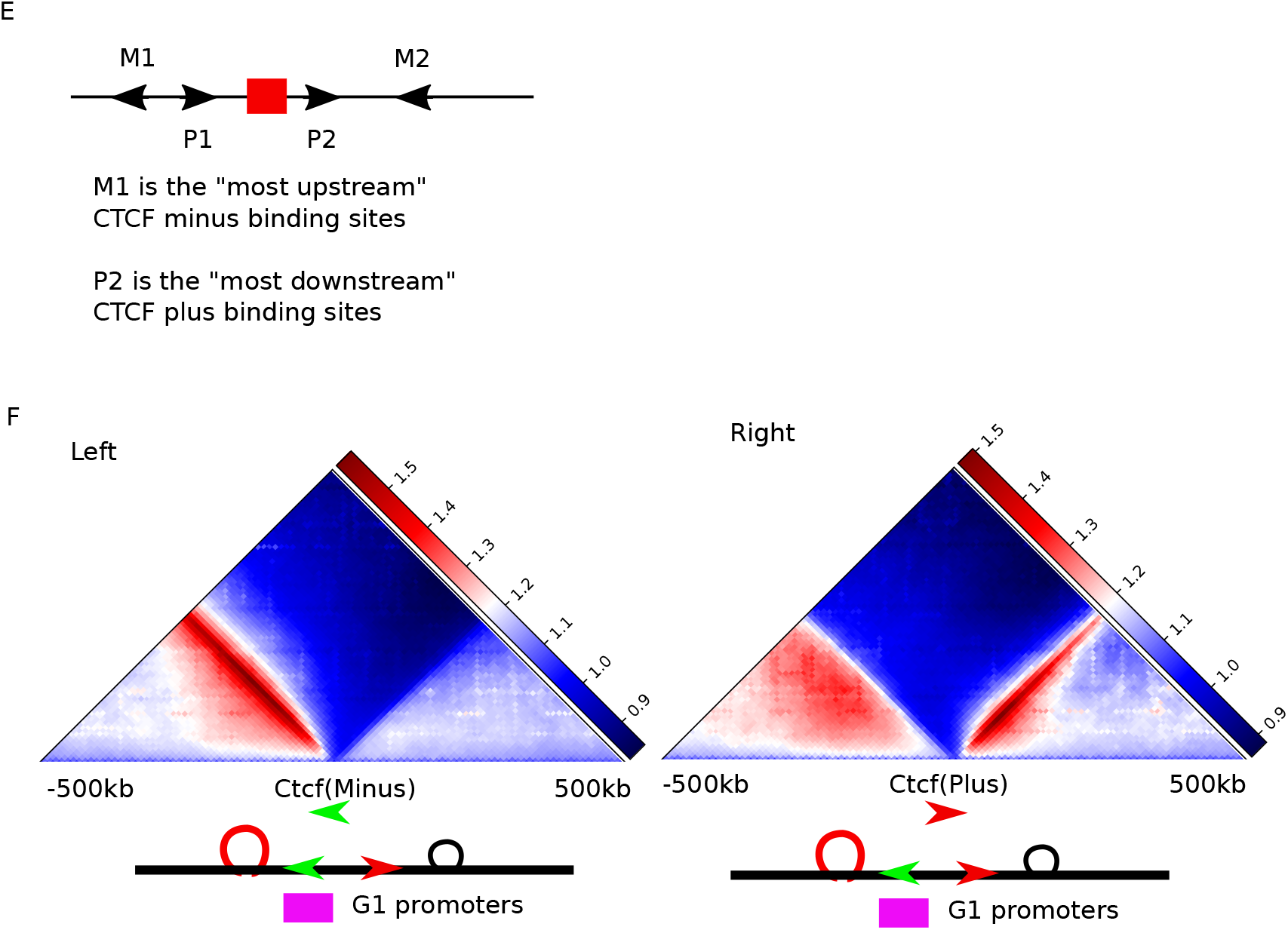
(A) The distribution of CBEP and CBEM relative to the G2 promoter. (B) The histogram shows the number of CBEM and CBEP in the 300kb regions flanking the G1 promoters. (C) Same as (B) but for the G2 promoters. (D) Same as the Fig6b but for G2 promtoers. Note that the high-intensity dots locate the upstream of the G2 promoters. (E) The two methods of choosing CBEP and CBEM for aggregation. The first method will choose M1 for CBEM aggregation and P2 for CBEP aggregation as they form the divergent pair with the largest distance. (F) The aggregate results for the CBEM (left) and CBEP (right). The CBEM and CBEP are selected according to the second methods (See Fig6c, which is based on the first method)

**Figure S6:**
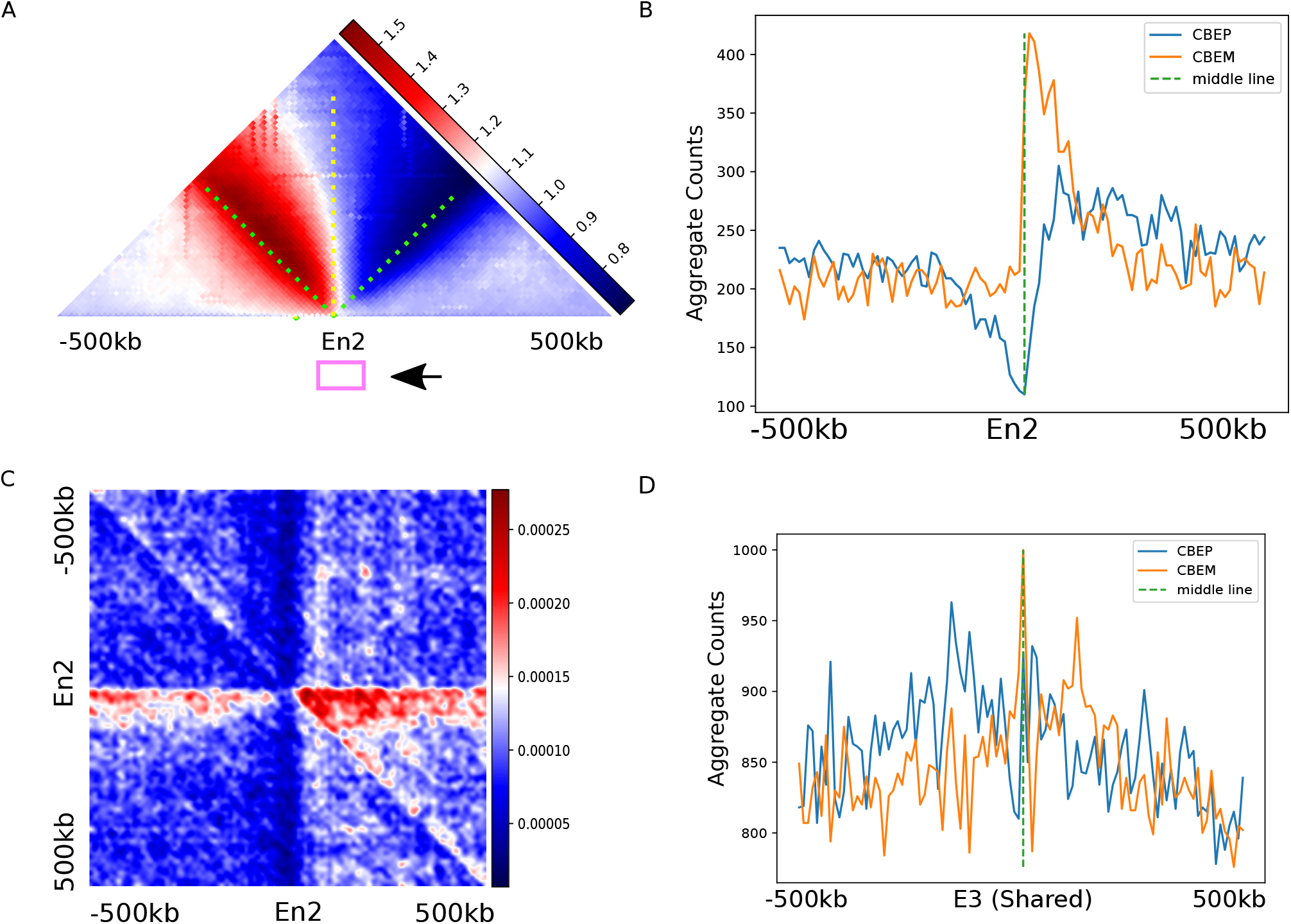
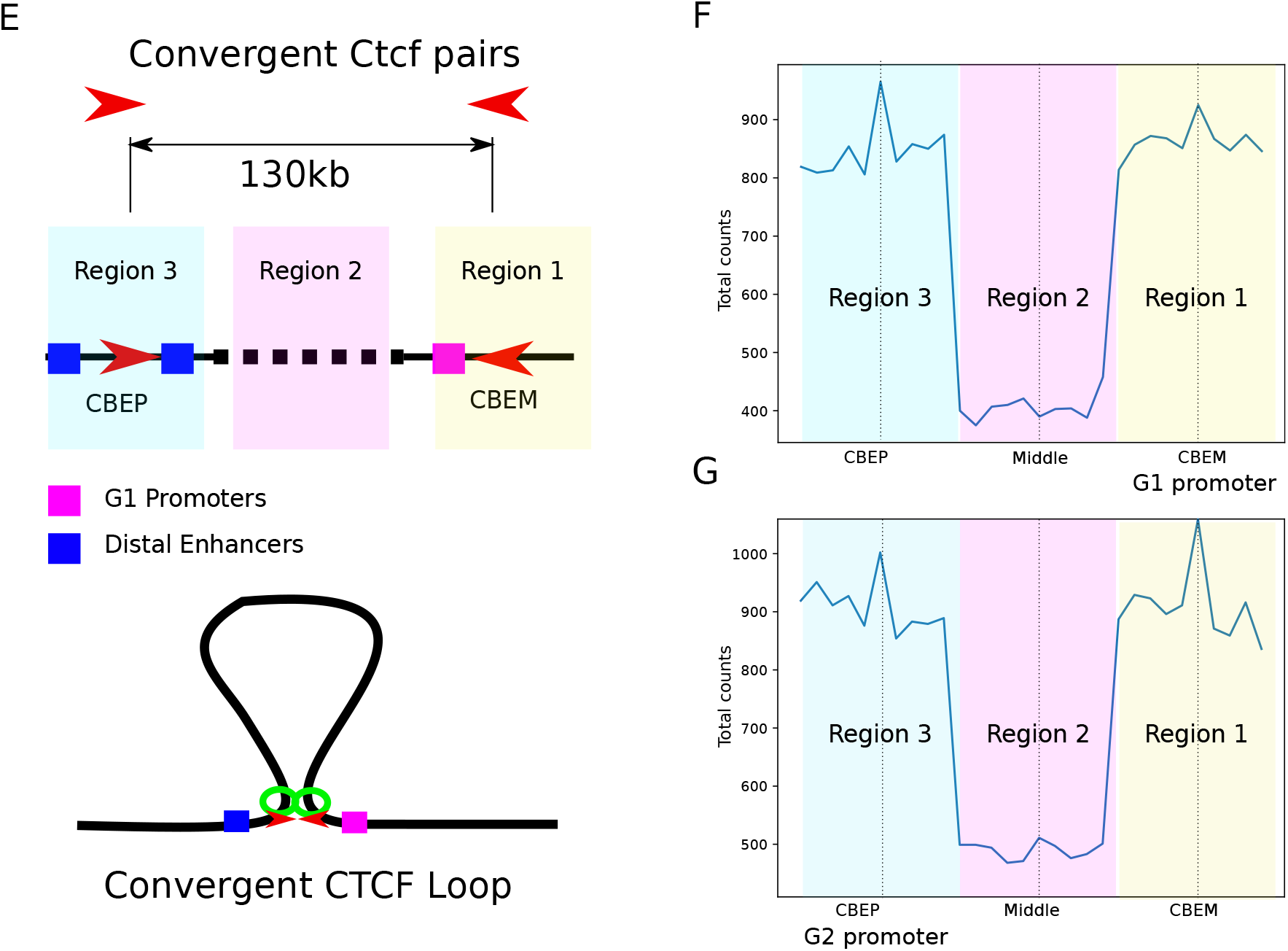
(A) The heatmap shows the E2AP, a pattern centered on the E2 enhancers. (B) The line plot shows the aggregate CBE signals centered on the E2 enhancers. (C) The joint probability heatmap shows that E2 enhancers surround the divergent CBE pairs. The divergent CBE pairs are located at the downstream side. (D) The line plot shows the distribution of CBE centered on the E3 enhancers. (E) The diagram shows the convergent CBE pairs are associated with the G1 promoters (magnet box). The median distance between the convergent CBE pairs is 130kb. Note that the G1 promoters and the associated CBEM are located within region 1. (F)The blue line shows the aggregate enhancer signal in three regions. The enhancers are more enriched within region 1 and 3 than that of region 2. Note that region 1 flanks the G1 promoter. (G) Same as (F) but for G2 promoters. Note that region 3 flanks the G2 promoter.

